# LETSmix: a spatially informed and learning-based domain adaptation method for cell-type deconvolution in spatial transcriptomics

**DOI:** 10.1101/2024.04.27.591425

**Authors:** Yangen Zhan, Yongbing Zhang, Zheqi Hu, Yifeng Wang, Zirui Zhu, Sijing Du, Xiangming Yan, Xiu Li

## Abstract

Spatial transcriptomics (ST) has revolutionized our understanding of gene expression patterns by incorporating spatial context. However, many ST technologies operate on heterogeneous cell mixtures due to limited spatial resolution. To resolve cell type composition at each sequencing spot, several deconvolution methods have been proposed. Yet, these approaches often underutilize spatial context inherent in ST data and paired histopathological images, meanwhile overlooking domain variances between ST and reference single-cell RNA sequencing (scRNA-seq) data. Here, we present LETSmix, a novel deconvolution method that enhances spatial correlations within ST data using a tailored LETS filter, and employs a mixup-augmented domain adaptation strategy to address domain shifts. The performance of LETSmix was validated across diverse ST platforms and tissue types, including 10x Visium human dorsolateral prefrontal cortex, ST human pancreatic ductal adenocarcinoma, 10x Visium mouse liver, and Stereo-seq mouse olfactory bulb datasets. Our findings demonstrate that the proposed method accurately estimates cell type proportions and effectively maps them to the expected regions, establishing a new record among current state-of-the-art models. LETSmix is expected to serve as a robust tool for advancing studies on cellular composition and spatial architecture in spatial transcriptomics.

**GRAPHICAL ABSTRACT:** 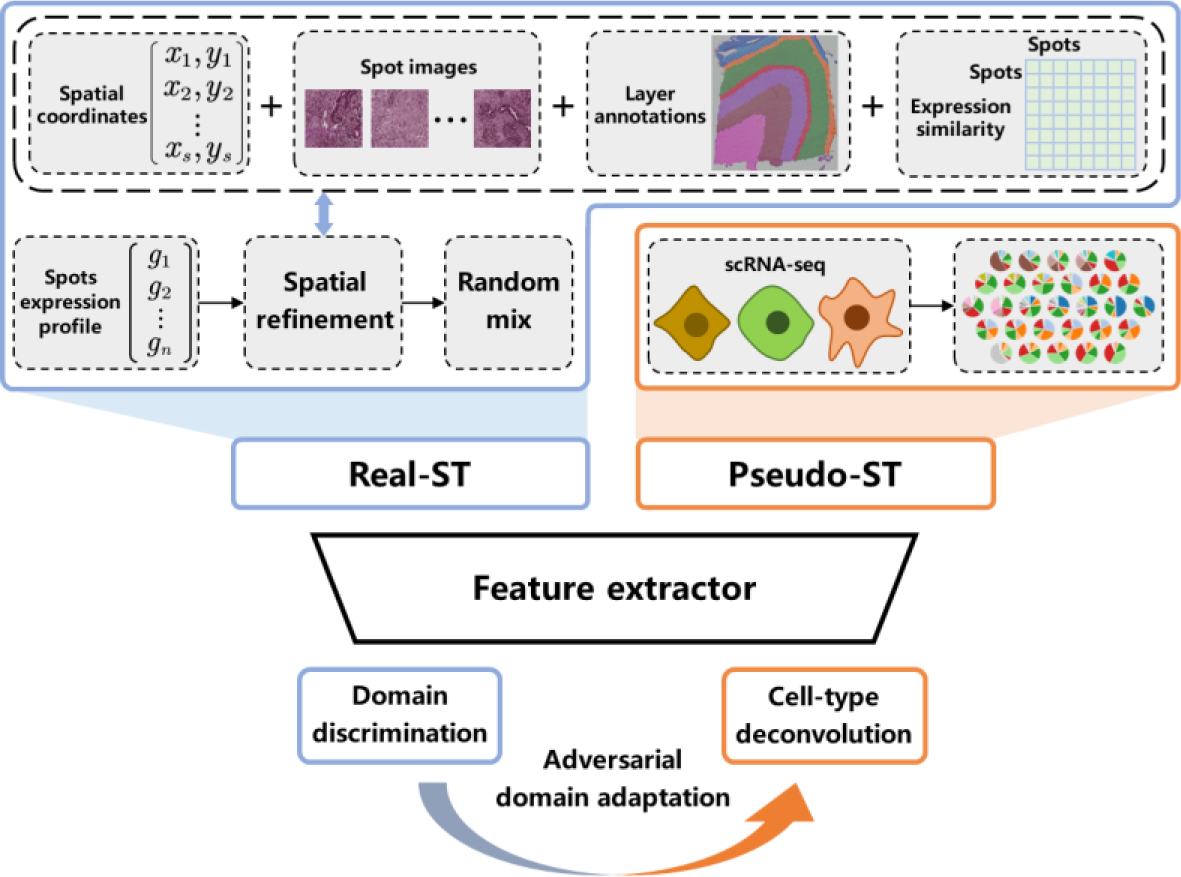

## INTRODUCTION

In the last decade, the continuous advancement of single-cell RNA sequencing (scRNA-seq) technology has facilitated high-throughput sequencing of individual cells, revealing comprehensive gene expression profiles within them (1,2). This technological leap has unearthed cellular heterogeneity, enabling the identification of diverse cell types, cell subpopulations, and transcriptional state alterations within complex cell populations (3). However, the intrinsic nature of scRNA-seq—sequencing individual cells—necessitates the isolation of cells from their native tissue prior to sequencing, hampering the integration of spatial information for analysis (4). To determine the spatial distribution of gene transcriptomes and the tissue microenvironment, spatial transcriptomics (ST) technology has emerged. This innovative approach enables the detection of gene expression profiles across numerous locations within tissue regions while retaining positional information (5,6). Currently, ST techniques are broadly categorized into image-based and sequence-based methods, which complement each other to some extent (7). Image-based methods such as MERFISH (8) and seqFISH+ (9) utilize fluorescent probes to target specific genes, offering high resolution even at the subcellular level (10). However, due to their reliance on targeted fluorescent probes, these techniques are confined to detecting a limited number of genes, typically a few hundred (11). In contrast, sequence-based techniques, such as Spatial Transcriptomics (ST) (12) and SLIDE-seq (13), can capture the expression profiles of a complete gene repertoire, but their sequencing resolution is lower, often encompassing multiple cells within each detection site, referred to as a “spot” (7). Notably, these sequence-based techniques usually provide not only the positional coordinates of each detection site but also H&E-stained histological images of the sequenced tissue. Despite ongoing advancements in improving their resolution, accurately assigning individual cells to each spot remains a challenge (14).

In sequence-based ST data, to discern the spatial distribution of diverse cell types and gain deeper insights into their compositional structure, prevailing methods integrate scRNA-seq with ST data (15,16). scRNA-seq provides gene expression profiles of individual cells alongside their corresponding cell type information, facilitating the inference of cell type-specific gene expression traits. The integration of this information with expression patterns observed in each spot from ST allows the analysis of contributions from each cell type at each spot (17-19). Current mainstream cell-type deconvolution methodologies can be broadly categorized into three types: statistical probabilistic models, matrix factorization-based models, and deep learning-based models (16,20,21). Probabilistic models such as RCTD (19), Stereoscope (22), Cell2location (23) and POLARIS (24) assume that gene expression in scRNA-seq and ST data follows certain distributions, such as the negative binomial or Poisson distribution. Although various influencing factors, such as technology sensitivity, batch effects, and per-location shifts, can be specifically parameterized within these methods, the regressed parameters may deviate from their original design. Similarly, these factors can also be transformed into parameter matrices in models based on nonnegative matrix factorization (NMF), as exemplified by SPOTlight (17), SpatialDWLS (18), and CARD (25). The learned cell-type gene signatures from scRNA-seq are used to disentangle each spot in the ST by the aforementioned two modeling strategies. On the other hand, deep learning approaches are gaining traction as promising solutions because of their capacity to apprehend and leverage intricate patterns and correlations within data. Techniques such as graph networks (26,27) capitalize on spatial relationships, while domain adaptation methods (28,29) bridge technical variances between scRNA-seq and ST. DSTG (30) constructs a graph network, generating synthetic data from scRNA-seq to approximate true cell-type proportions in ST. Furthermore, self-supervised training with variational graph autoencoders applied in Spatial-ID (31), tissue histological image integration applied in SpaDecon (32), and domain-adversarial learning applied in CellDART (33) are innovative deep learning strategies that improve cell-type deconvolution accuracy in ST data.

However, existing methods face two significant limitations: insufficient utilization of spatial contextual information and inadequate consideration of domain differences between ST and scRNA-seq data. First, many approaches fail to fully leverage the rich spatial information available in ST datasets, such as spatial coordinates, histological images, and region-specific annotations. Second, these methods rarely account for the domain discrepancies between ST and scRNA-seq data, which arise from differences in sequencing technology, gene detection sensitivity, and sample preparation. For instance, while SpaDecon incorporates spot coordinates and histological image data, it is directly applied to ST deconvolution after being trained on scRNA-seq data, overlooking the domain variance and other critical spatial features like region annotations. Similarly, although CellDART employs domain adaptation techniques, it fails to consider potential spatial correlations and does not address the significant sample size imbalance between the source (scRNA-seq) and target (ST) domains, leading to suboptimal performance in real-world applications.

In this work, we propose LETSmix, the first method, to our knowledge, that simultaneously addresses both challenges by leveraging spatial context information and incorporating domain adaptation. Furthermore, our method introduces several key innovations in both areas. With respect to spatial information, LETSmix goes beyond previous methods like SpaDecon by incorporating region annotations and gene expression similarity, and constructing a LETS filter to capture more finegrained spatial correlations between spots in ST data. To address domain differences, LETSmix extends the domain adaptation strategy used in CellDART by introducing a novel mixup-based data augmentation process to mitigate the issue of sample size imbalance between the source and target domains, and optimizing the training procedure to enhance the stability of the domain adaptation learning process. Together, these innovations allow LETSmix to more effectively align scRNA-seq and ST data, ensuring robust and accurate cell-type deconvolution across domains.

Furthermore, we conducted comprehensive experiments to validate LETSmix on multiple datasets derived from different ST technologies and tissues. Each dataset presents unique characteristics, such as the well-defined layer structure in the 10x Visium human dorsolateral prefrontal cortex dataset (34), the inclusion of both internal and external data and the limited number of spots in the ST human pancreatic ductal adenocarcinoma dataset (35), the dominance of hepatocytes in the 10x Visium mouse liver dataset (36), which poses challenges for identifying rare cell types, and the single-cell spatial resolution of the Stereo-seq mouse olfactory bulb dataset (37), which allows us to evaluate the applicability of LETSmix to cutting-edge ST technologies. Across all these datasets, the proposed method consistently outperforms state-of-the-art methods, demonstrating its versatility and superior performance under diverse biological conditions.

## MATERIAL AND METHODS

### Public dataset collection

*Human dorsolateral prefrontal cortex (DLPFC) data.* The 10X Visium DLPFC dataset was derived from a postmortem 30-year-old neurotypical subject (34). Our experiments included all twelve ST samples, each containing 3000-5000 spots with 33,538 common genes. Layer annotations were provided by assigning each spot to a specific cortical layer, ranging from L1 to L6 and WM (white matter), based on layer-specific marker genes and expert inspections (38). A few spots without layer annotations were excluded in our experiments. The reference single-nucleus dataset (also referred to as scRNA-seq) was obtained from the DLPFC tissues of different postmortem individuals without neurological disorders (39). This scRNA-seq dataset contained 56,561 cells and 30,062 genes. We focused on 10 layer-specific excitatory neurons to evaluate different cell-type deconvolution models using AUC and ER metrics.

*Human pancreatic ductal adenocarcinoma (PDAC) data.* The PDAC dataset was acquired from two tumorous tissue sections of two patients, denoted as PDAC-A and PDAC-B (35). The ST datasets were collected using the Spatial Transcriptomics technology, while their matched scRNA-seq datasets were obtained through inDrop. The paired ST and scRNA-seq data were both derived from the same tissues of the same patients. Additionally, an external scRNA-seq dataset named PDAC-Peng (40) obtained through 10x Chromium was used to evaluate model performance under unmatched conditions. Cell types and their compositions differs among the three scRNA-seq datasets. Although the true cell type composition of each spot in the ST data of PDAC-A and PDAC-B remains unknown, the overall composition of the whole tissue, i.e., the average cell-type proportions in all spots, is expected to be close to that of the matched reference scRNA-seq dataset. Thus, a JSD value can be calculated to measure the consistency between the estimated overall cell type composition and the expected ground truth. Moreover, a few annotated cell types in the reference scRNA-seq dataset are assumed to be enriched within specific tissue regions (Supplementary Table S1). Based on this prior knowledge, the ER metric was implemented to compare the performances of different deconvolution methods in predicting the regional distribution patterns of these cell types.

*Mouse liver (Liver) data.* The Liver dataset was obtained from a published study (36). Three consecutive Visium slices of healthy mouse liver tissues were analyzed in this study, each delineated into five distinct regions. These ST samples are primarily composed of hepatocytes, making it challenging for deconvolution tools to accurately identify other cell types. The reference scRNA-seq datasets includes cells obtained from three different digestion protocols: ex vivo digestion, in vivo liver perfusion, and frozen liver single-nucleus RNA-seq (referred to as “ex vivo scRNA-seq”, “in vivo scRNA-seq”, and “nuclei scRNA-seq”, respectively) (21). All three scRNA-seq datasets contain the same nine cell types. Notably, portal vein and central vein endothelial cells (ECs) are expected to be exclusively present in the portal and central regions, respectively. Moreover, previous studies based on confocal microscopy have suggested that the average proportion of each cell type in all spots within the ST sample is equivalent to that in the nuclei scRNA-seq dataset (21,36). Therefore, JSD values were computed to evaluate the estimated proportions in ST. Cell-type proportions in the ex vivo and in vivo scRNA-seq datasets were adjusted to match those in the nuclei scRNA-seq dataset.

*Mouse olfactory bulb (MOB) data.* The ST dataset was collected using Stereo-seq (37), which is an emerging spatial omics platform with subcellular spatial resolution. Here, it was binned to a cellular-level resolution (∼14 μm), allowing for more manageable data analysis. Seven layers of the laminar organization in MOB were annotated by Xu et al. (27), including the rostral migratory stream (RMS), granule cell layer (GCL), internal plexiform layer (IPL), mitral cell layer (MCL), external plexiform layer (EPL), and olfactory nerve layer (ONL). Given the large number of spots in this ST dataset (approximately 20,000), we randomly selected half of the spots to expedite computation. For cell-type deconvolution, a publicly available scRNA-seq dataset generated using 10x Chromium from the same tissue source was used as the reference (41), which originally contained 38 cell types. In our experments, we merged several cell subtypes, resulting in 27 distinct cell types used for model training. To evaluate the deconvolution performance, we employed the ER metric to assess the enrichment of specific cell types in their expected regions. Additionally, due to the well-characterized laminar structure of the olfactory bulb, which presents distinct inside-out regional patterns, we used Moran’s I to analyze the spatial autocorrelation of the deconvolution results, providing further insights into the spatial organization of the predicted cell types.

### Implementation of LETSmix

A schematic diagram illustrating the proposed method is presented in Fig. 1. The overall network framework comprises three main components. First, an adjacency matrix termed as “LETS filter” was constructed leveraging information from Layer annotations, gene Expression similarities, histological image Texture features, and Spot coordinates to accurately capture the spatial correlations among different spots. This matrix was subsequently employed to perform local smoothing on the ST dataset, emphasizing spatial relationships between neighboring spots with similar morphological features. Second, a fixed number of cells were randomly selected from the scRNA-seq dataset to synthesize gene expression data for each pseudo-spot. Meanwhile, the spatially refined real-ST data were also randomly mixed to generate more samples in the target domain. In the final stage, the synthesized pseudo-ST data and mixed real-ST data were simultaneously fed into a deep learning network for adversarial domain adaptation training. Here, the domain classifier aims to differentiate the domain of the input spot expression data, while the source classifier estimates cell type compositions within each spot in both the pseudo-ST and real-ST datasets.

**Figure 1:**
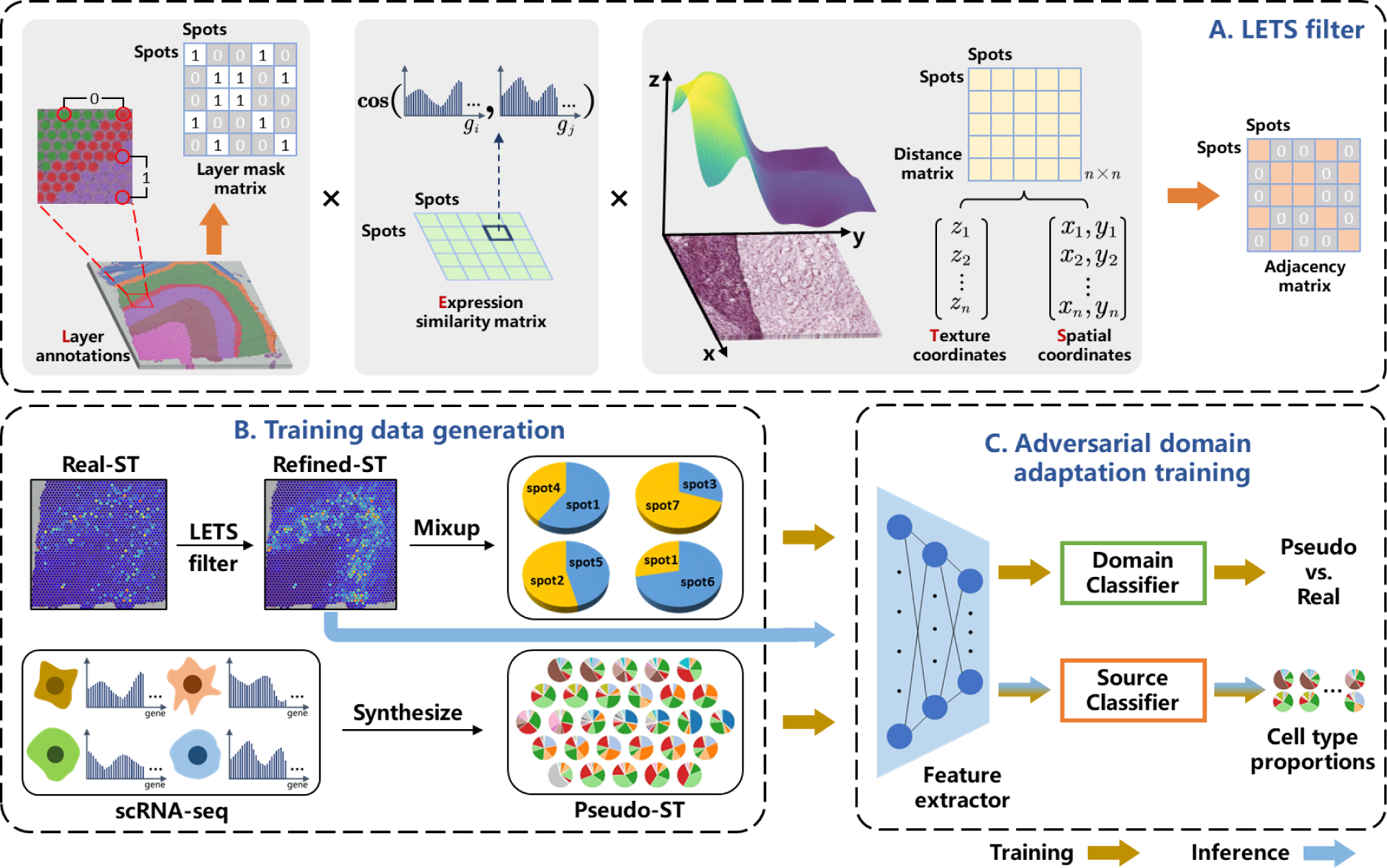
Schematic overview of the LETSmix workflow. LETSmix is designed to perform cell-type deconvolution for spatial transcriptomics based on the labeled reference scRNA-seq data. (A) An adjacency matrix reflecting internal correlations between spots in ST was constructed leveraging information from layer annotations, spot gene expression, histological image texture features, and spot spatial coordinates. (B) Pseudo-ST data with known cell type compositions were synthesized by randomly selecting cells from the reference scRNA-seq dataset. The real-ST data were refined by the LETS filter constructed from (A) and then underwent a mixup procedure for data augmentation. (C) The network structure comprises a shared feature extractor and two classifiers: the source classifier estimates cell type proportions, and the domain classifier identifies spots as real or pseudo. The two branches are trained in an adversarial manner, aiming to eliminate domain shifts in the extracted features. After training the model, the spatially refined real-ST data were directly fed into the feature extractor and the source classifier for cell-type deconvolution.

*Spatial refinement of the ST data.* The expression data can be noisy due to the limited number of cells in each spot. To address this issue, we constructed an adjacency matrix that aggregates information from neighboring spots. Initially, the distance *d*_*i*,*j*_ between spots *i* and *j* was calculated by incorporating spatial, morphological, and expression features. Following the method adopted in SpaDecon, the coordinate *z*_*i*_ for spot *i*, representing its histological image texture features, is defined as

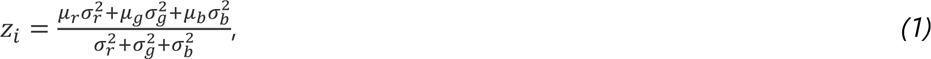

where *μ*_*r*_, *μ*_*g*_, *μ*_*b*_ are the means of the RGB values across all pixels in the ℎ × ℎ image patch of spot *i*, and ℎ is the diameter of each spot. *σ*_*r*_, *σ*_*g*_, *σ*_*b*_ are the standard deviations of *μ*_*r*_, *μ*_*g*_, *μ*_*b*_, respectively, across all spots. Then, *z*_*i*_ was further scaled as

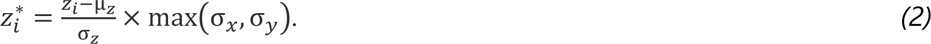

Here, μ_*z*_ and σ_*x*_, σ_*y*_, σ_*z*_ are the means and standard deviations, respectively, of the (*x*, *y*, *z*) coordinates of all the spots in the ST data. This scaling ensures that the weight of the histological features approximately matches that of the spatial location features. The Euclidean distance between two spots *i* and *j* using the three coordinates is given by

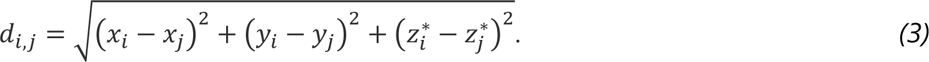

The total number of spots in the real-ST data is denoted by *n*, and the adjacency matrix A = [*a*_*i*,*j*_]_*n*×*n*_ is constructed based on *d*_*i*,*j*_ combined with expression data and layer annotation information, as

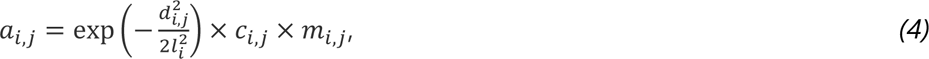

where *l*_*i*_ is a scaling factor controlling the degree of local smoothing, *c*_*i*,*j*_ is the cosine similarity between the expression data of two spots, and *m*_*i*,*j*_ is an element from a mask matrix M = [*m_i,j_*]_*n*×*n*_ constructed using layer annotations, as

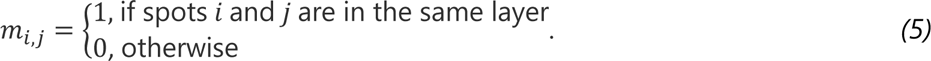

Note that SpaDecon only utilizes spatial coordinates and histological images to construct the adjacency matrix, whereas we leverage additional information from layer annotations and gene expression data. This improvement introduced by our designed adjacency matrix accurately reflects the intrinsic spatial correlations within the ST data, thereby enhancing the accuracy of the cell-type deconvolution results. Next, we describe the selection strategy of an appropriate value for *l*_*i*_. Define *s*_*i*_ by

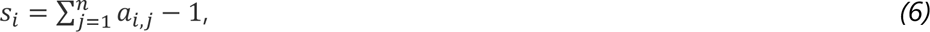

which represents the summed weights of other spots with respect to the target smoothing spot *i*. The value of *s*_*i*_ is determined by *l*_*i*_. Although the relationship between them cannot be definitively ascertained through numerical methods, it is established that *s*_*i*_ increases monotonically with respect to *l*_*i*_. Thus, we start by specifying the desired value of *s*_*i*_ and then iteratively approach the value of *l*_*i*_, as outlined in Supplementary Algorithm S1. In our experiments, *s*_*i*_ of different spots where *i* ∈ {1,2, … , *n*} were set to be the same value *s̃*. Note that SpaDecon employs a similar approximation technique for determining the value of *l*. However, the scaling factor *l* in SpaDecon is a scalar, ensuring that only the average summed weights of other spots are a specific value, resulting in varying degrees of local smoothing for different spots. Although there are *n* different values in the vector *l* to be approximated in our improved method, the computational complexity of the designed algorithm is similar to that of the original algorithm. It requires only a single round of approximation to directly compute each value in the vector *l*, thereby not compromising the operational speed of the model. With the constructed adjacency matrix *A*, we performed matrix multiplication to obtain smoothed gene expression data *G_r_*^∗^ of real-ST data, as follows:

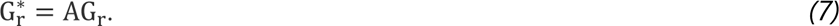

*Mixup of the ST and scRNA-seq data.* The expression data for each pseudo-spot were synthesized from randomly selected *k* cells in the reference scRNA-seq dataset. We used [*g*_*c*1_, *g*_*c*2_, … , *g*_*ck*_] to represent the gene expression profile of each selected cell. Moreover, random weights [*w*_*c*1_, *w*_*c*2_, … , *w*_*ck*_] were assigned to each cell, representing its proportion in the generated pseudo-spot, and the sum of the *k* weights was constrained to 1. Thus, the gene expression profile of the pseudo-spot is the weighted sum of *k* cells, denoted as:

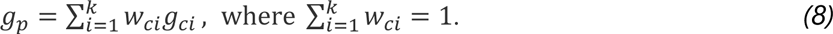

Similarly, the ground truth cell-type proportions in this pseudo-spot are also the weighted sum of the cell types of the selected *k* cells. In this way, the total amount of gene expression in a pseudo-spot is equivalent to that in a cell according to the scRNA-seq data.

Through various combinations, an infinite number of pseudo-spots can be synthesized, whereas the quantity of real-spots is typically limited. To address the discrepancy in data volume between the target domain and the source domain during the subsequent domain adaptation process, we also applied mixup to real-ST data refined by the LETS filter, thereby significantly increasing the amount of data in the target domain. Specifically, a mixup ratio λ was randomly sampled from a Beta distribution *Beta*(α, α) for α > 0. The resulting mixed-spot was generated by:

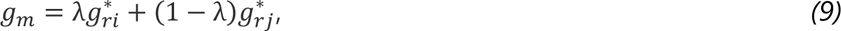

where *g*^∗^*_ri_* and *g*^∗^*_rj_* are two randomly selected refined real-spots. In fact, mixup is a commonly used data augmentation strategy in computer vision. We find it particularly suitable for ST data, where concerns about its potential negative impacts such as the loss of target images or confusing merged objects are not as relevant.

*Adversarial domain adaptation training for the deep learning network.* The training process of the LETSmix network, which consists of nonlinearly activated fully connected layers (detailed in Supplementary Table S2), was conducted using the PyTorch-gpu package (version 1.12.0). Initially, a shared feature extractor *F* was utilized to derive low-dimensional features from both pseudo-ST and locally smoothed real-ST data. The subsequent source classifier *S* and domain classifier *D* were trained in an adversarial manner. Specifically, we first pretrained the feature extractor and the source classifier using only the pseudo-ST data for cell-type deconvolution, leaving the domain classifier aside. Then, the two branches were trained alternatively. In the domain classifier training procedure, the learnable weight parameters in the feature extractor and source classifier were frozen. This process aims to effectively distinguish whether the extracted features originate from pseudo-ST or real-ST. At this stage, the loss function is defined as the binary cross-entropy between the predictions of the domain classifier and the assigned domain label, formalized as

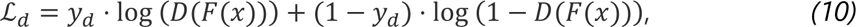

where *x* and *y*_*d*_ denote the input data and the domain label (0 for pseudo-ST and 1 for real-ST), respectively. In the source classifier training procedure, we froze the trained domain classifier and used it to classify the domain of the input data. At this stage, the feature extractor aims to deliberately confuse the domain discriminator, preventing it from accurately distinguishing the domain of the extracted features. This serves the purpose of eradicating domain variances existing in the feature space. Simultaneously, the source classifier endeavours to accurately estimate the proportions of different cell types within each input pseudo-spot. Thus, there are two components in the loss function. The first component is defined as the Kullback-Leibler divergence (KLD) between the estimations of the source classifier and the cell-type proportion labels, while the second component is the binary cross-entropy between the predictions of the frozen domain classifier and the inverted assigned domain label, which can be formalized as

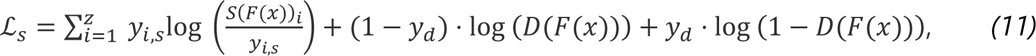

Here, *z* is the total number of cell types, while *y*_*i*,*s*_ and *S*(*F*(*x*))_*i*_ stand for the label and estimation for the *i*th cell type, respectively. Note that the KLD loss is only calculated for input data from the source domain (pseudo-ST), while the binary cross-entropy loss is calculated for data from both domains. Maintaining the same batch size for data from pseudo-ST and real-ST, we find that it is crucial for the training iterations of the domain classifier to be more than that of the source classifier within a cycle of alternation, where the ratio between the two branches is *d*: 1. This observation aligns with logical expectations because the ability of the domain discriminator to guide the feature extractor in learning domain-invariant features hinges upon its accurate domain differentiation.

### Preprocessing spatial and single-cell datasets

The preprocessing of the ST and scRNA-seq data was conducted in Python using the Scanpy package (version 1.9.3). Initially, *m* highly expressed marker genes for each cell type in the scRNA-seq data were selected through the Wilcoxon rank-sum test. Note that the expression data from the scRNA-seq data were normalized and log1p-transformed before the marker gene selection process, ensuring an equivalent total amount of gene expression for each cell. The subsequent procedures utilized the intersection between the extracted marker genes and all genes in the ST data. Then, the expression data of the selected marker genes from both the scRNA-seq and ST data were once again normalized and log1p-transformed. Importantly, the normalized total amount of expression in each cell from the scRNA-seq data should be (1 + *s̃*) times greater than that in each spot from the ST data. This adjustment accounts for the real-ST refinement process, ensuring that the total counts in a generated pseudo-spot and a real spot are equal. Finally, the expression data in pseudo-ST and real-ST were min-max scaled, to ensure that the values lie between 0 and 1. This comprehensive preprocessing pipeline sets the stage for subsequent analyses in LETSmix, enhancing the comparability and accuracy of the deconvolution results across different datasets.

### Evaluation metrics

In the experiments presented in this paper, four quantitative metrics were employed to evaluate the performance of various cell-type deconvolution methods: Area Under the Curve (AUC), EnRichment (ER), Jensen-Shannon Divergence (JSD) and Moran’s I. The AUC and ER gauge the agreement between the predicted spatial distribution patterns of regionally restricted cell types and layer annotations, while JSD assesses the alignment of the estimated general cell type composition in the ST dataset with prior knowledge from the matched scRNA-seq dataset. Moran’s I, on the other hand, is used to assess the spatial autocorrelation of the predicted cell-type distributions, specifically evaluating the similarity of cell type compositions within spatially adjacent spots. A high Moran’s I value indicates strong spatial clustering of similar cell types, which is expected in certain tissues with clear structural organization. In the context of receiver operating characteristic (ROC) analysis, a spot is labeled 1 if it resides in the target region of the layer-specific cell type under examination; otherwise, it is labeled 0. The AUC value is then calculated between the estimated proportions of this cell type in all spots and the assigned labels. Despite the common use of the AUC metric in other cell-type deconvolution studies (33,42), it is acknowledged to have certain limitations. Notably, not all spots within the target region necessarily contain the layer-specific cell type under investigation in reality. Additionally, setting the label to 1 does not realistically represent the actual proportion of this cell type in a spot within the target region. Therefore, even if the model accurately predicts cell type compositions in each spot, achieving an AUC value of 1 is improbable. To address these limitations, an additional metric called EnRichment (ER) was introduced in this study. The ER calculates the estimated total counts of a layer-specific cell type within the target region divided by the total counts of that cell type in the entire tissue section. Due to the unknown total number of cells in each spot, this value was approximated using the estimated proportions of that cell type as a substitute. The formalization of the ER is given by:

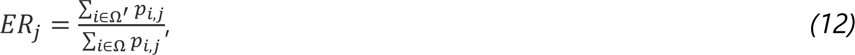

where Ω^′^ and Ω denote the spots in the target region and the entire tissue section, respectively. *p*_*i*,*j*_ is the estimated proportion of the *j*th cell type in the *i*th spot. In cases where multiple cell types share a common target region, such as cancer clone A and cancer clone B cells in the PDAC-A dataset, their estimated proportions in each spot are first summed together as a broader cell type, and an ER value is calculated based on this merged cell type. To obtain JSD values, we first computed the overall proportions of various cell types in both the scRNA-seq and ST data. The calculation for the latter involved averaging the cell type compositions across all estimated spots, as illustrated by Formula (13).

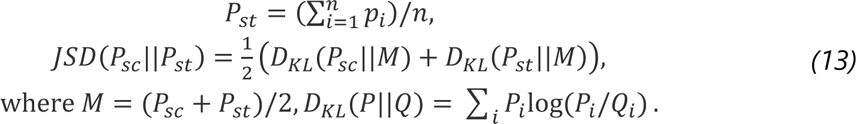

All the quantitative results presented in this paper were obtained from five repeated experiments. The models under evaluation were trained from scratch five times with different random seeds. In each repeated experiment, the learned parameters in LETSmix were saved at the training epoch where the best performance was achieved. The distribution patterns of 10 layer-specific cell types in the DLPFC datasets estimated by different models were examined using the AUC and ER metrics. The AUC metric was also utilized in the ablation and hyperparameter analysis for the DLPFC dataset. For the PDAC dataset, ER and JSD values were calculated for different models under matched conditions, and only the ER values were computed under unmatched conditions. Similarly, the ER and JSD metrics were applied to the Liver dataset to analyze the estimated distribution patterns of two regionally restricted cell types and the general cell type compositions, respectively. In the MOB dataset, ER and Moran’s I values were computed for certain cell types with potentially regional distribution patterns.

### Method comparison

The deconvolution performance of LETSmix was benchmarked against that of five other state-of-the-art computational methods: CellDART (33), SpaDecon (32), Cell2location (23), POLARIS (24), and CARD (25). CellDART and SpaDecon are two deep learning-based models, and can be seen as different simplified versions of LETSmix. CellDART also follows a source-domain dual-branch framework, leveraging the adversarial domain adaptation strategy to bridge technical variances between ST and the scRNA-seq data, but completely ignores the inherent correlations among spots in the ST dataset and the significant imbalance between the number of samples in the source and the target domain. SpaDecon simply utilizes information from spot coordinates and histological images to locally smooth the ST dataset. The model is trained on scRNA-seq data with a supervised clustering algorithm, and is directly employed to infer cell-type proportions in ST. Cell2location assumes that gene expression data in scRNA-seq and ST data follow a negative binomial distribution. Cell type signatures were inferred from the reference scRNA-seq dataset and used to estimate cell type proportions decomposed from the ST data. The model incorporates parameters capturing various sources of data variability. POLARIS, like Cell2location, is also a statistical probabilistic model. While it accounts for fewer sources of variability than Cell2location, POLARIS uniquely incorporates region annotations, allowing for the possibility that gene expression levels may vary across different regions within the tissue. In CARD, the count matrices of the scRNA-seq and ST data are decomposed into different parameter matrices. Cell-type signatures were inferred from the reference scRNA-seq dataset, and the cell type composition matrix was estimated from the ST data via NMF regression. The model introduces a conditional autoregressive assumption, leveraging position coordinate information to consider spatial correlations among spots. For all methods, the default parameters were applied for analyses unless otherwise specified.

### Parameter selection

The key parameters of the proposed LETSmix method include the degree of spatial context information used (*s̃*), the ratio of training iterations between the domain classifier and the source classifier (*d*), the number of marker genes per cell type (*m*), and the number of cells per pseudo-spot (*k*). To determine the optimal values for these parameters, we investigated the performance variation of LETSmix in the DLPFC dataset across a spectrum of conditions as a reference (Supplementary Figure S1).

Different choices for *s̃* revealed that optimal performance was achieved when its value ranges from 0.5 to 1. Extremes in the utilization of spatial background information, either too low or too high, impeded the cell-type deconvolution process. Considering that a parameter similar to *s̃* is incorporated in SpaDecon, we set the value of *s̃* to 0.5, which is the default value applied in SpaDecon, for a fair comparison.

Effective discrimination by the domain discriminator between the two domains is crucial in guiding the feature extractor to eliminate domain differences, which is substantiated by the fact that more training iterations for the domain discriminator led to enhanced performance. However, improvements in deconvolution performance stopped when *d* reaches 10, as the discriminative capacity of the domain discriminator itself has inherent limitations. Notably, the official implementation in CellDART does not prioritize additional training iterations for the domain classifier. In our reproduced experimental outcomes, adhering to the training methodology outlined in the public code of CellDART yielded notably lower accuracy in deconvolution results compared to those reported in the original paper. To maintain fairness, all experiments in this paper maintained a proportional ratio of 10:1 for training iterations between the domain and source classifiers within both LETSmix and CellDART.

Increasing the number of marker genes per cell type during the data preprocessing stage consistently yielded improved results, providing more comprehensive information for the model but at the expense of greater computational cost. Empirically, the value of *m* was set to 50 for all three public datasets used in this study.

Similar to *m*, increasing the value of *k* consistently led to enhanced deconvolution performance. With sufficient data, the model can learn more diverse features and adapt to complex application scenarios by including more cells in each pseudo-spot. In our experiments, *k* was set to 8 for all datasets because it is the default value applied in CellDART for a fair comparison.

## RESULTS

### Overview of LETSmix

The proposed cell-type deconvolution method integrates spatial correlations among spots while eliminating domain variances between the ST and the reference scRNA-seq data. As illustrated in Figure 1, both the scRNA-seq and ST data were preprocessed before they were fed into the deep learning network. The synthesis of pseudo-ST from annotated scRNA-seq data was implemented by randomly selecting cells to generate pseudo-spots, with their total gene expression counts determined by a weighted sum of the chosen cells. This procedure accumulated a substantial corpus of pseudo-ST data with known cell type compositions, facilitating supervised training of the feature extractor and source classifier. Real-ST data were locally smoothed with ancillary spatial context information. Considering potential noise in expression data due to limited cell numbers per spot and sequencing techniques, it is rational to aggregate information from neighboring spots with similar histological and gene expression features (32). To this end, an adjacency matrix, termed the LETS filter, was designed to signify spatial, histological, and gene expression similarities among real-ST spots. Further optimization of this matrix incorporated layer annotations as masks, restricting information sharing to spots within the same layer. Multiplying the constructed adjacency matrix with the original real-ST expression count matrix effectively enhances the data quality. Due to inherent technical differences between ST and scRNA-seq, the refined real-ST and synthetic pseudo-ST data underwent adversarial training within the deep learning network, employing a label inversion technique commonly used in domain adaptation methods. This technique ensures that the learned domain classifier effectively distinguishes real-spots from pseudo-spots, while the extracted features try to deceive the trained domain classifier. Notably, augmented training data were generated by mixing spots randomly selected from the spatially refined real-ST data to compensate for the significant disparity in data volume between real-spots and pseudo-spots. Thus, the source classifier, trained exclusively with labeled pseudo-ST, demonstrates superior performance in estimating cell type compositions from real-ST data.

To assess the efficacy of our proposed deconvolution method, LETSmix was applied to four representative public real ST datasets (Supplementary Table S3). These datasets encompass 12 ST samples of human brain cortex (DLPFC) data, 2 ST samples of human pancreatic ductal adenocarcinoma (PDAC) data, 3 ST samples of mouse liver (Liver) data, and 1 ST sample of Mouse olfactory bulb (MOB) data. All the ST samples in the DLPFC dataset share a common reference scRNA-seq dataset, while each PDAC ST sample is paired with a matched reference scRNA-seq dataset, supplemented by an external scRNA-seq dataset. The Liver dataset comprises 3 distinct scRNA-seq datasets with different sequencing protocols, and only one scRNA-seq dataset is used to deconvolve MOB ST data. The performance of LETSmix was benchmarked against that of other state-of-the-art methods (Supplementary Table S4), including CellDART, SpaDecon, Cell2location, POLARIS, and CARD, through qualitative heuristic inspection and quantitative evaluation using metrics such as Area Under the Curve (AUC), EnRichment (ER), Jensen-Shannon Divergence (JSD), and Moran’s I. Among these metrics, the AUC and ER evaluate the enrichment of regionally restricted cell types within target areas, while the JSD measures the consistency of the overall cell-type proportions between the entire ST tissue region and the matched reference scRNA-seq dataset. Moran’s I assesses the spatial autocorrelation of cell type distributions, indicating whether specific cell types exhibit clustered or dispersed spatial patterns.

### Benchmarking and robustness evaluation of LETSmix in deconvolution of human DLPFC data

We assessed the performance of LETSmix using a 10X Visium dataset derived from postmortem neurotypical human DLPFC tissues (35). In this dataset, 12 ST samples exhibit clear structural stratification, ranging from L1 to L6 and WM (white matter) annotated by pathologists (Figure 2A, Supplementary Figure S2). A shared reference scRNA-seq dataset (37) was applied to train each deconvolution method for benchmarking, as visualized by the UMAP representation in Figure 2B. Among the 28 annotated cell types, the spatial mapping results of 10 layer-specific excitatory neurons were used to quantify the deconvolution performance of the different tools.

**Figure 2:**
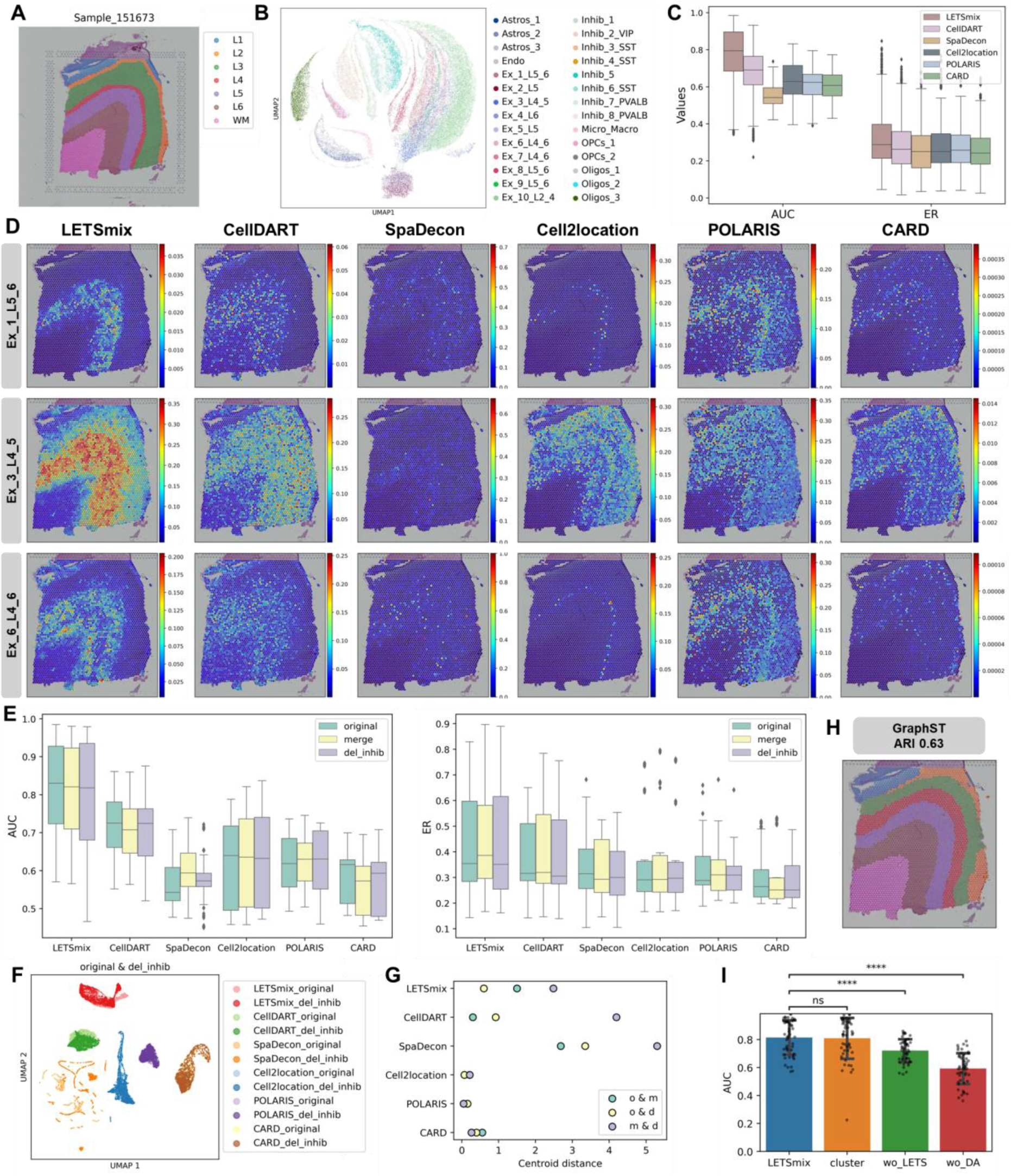
Application to the human brain cortex 10X Visium dataset. (**A**) Layer annotations of the ST sample named “151673” in the DLPFC dataset. (**B**) UMAP representation of the reference scRNA-seq dataset. (**C**) Box plots displaying the calculated AUC and ER values for the estimated cell type distribution in all 12 ST samples. Each box comprises 50 datapoints that represent scores for 10 layer-specific cell types in 5 repeated experiments, and ranges from the third and first quartiles with the median as the horizontal line, while whiskers represent 1.5 times the interquartile range from the lower and upper bounds of the box. (**D**) Estimated proportion heatmaps of 3 layer-specific excitatory neurons by each deconvolution method. (**E**) Box plots showing the calculated AUC and ER values for the estimated cell type distribution in the “151673” ST sample. “original” includes all 28 cell types in the scRNA-seq dataset. “merge” indicates that the cell subtypes were merged before model training except for the 10 excitatory neurons. “del Inhib” indicates that inhibitory neurons were deleted from the scRNA-seq dataset. (**F**) UMAP representation of deconvolution results from different methods under the “original” and “del Inhib” conditions. (**G**) Scatter plots of cluster centroid distances in the deconvolution results UMAP computed for each method under different condition pairs. (**H**) Clustering results of the “151673” ST sample given by GraphST. (**I**) Ablation study on the “151673” ST sample. “cluster” represents the situation where LETSmix leverages clustering results given by GraphST as the layer annotation information. “wo_LETS” represents the situation where LETSmix ignores all spatial context information, and “wo_DA” represents the situation where LETSmix is trained without the implementation of the domain adaptation strategy. Error bars represent the mean ± standard deviation. An independent t-test was performed between LETSmix and the other ablated models. Statistical significance is indicated above the bars (ns: not significant, ****P-value < 0.0001).

Figure 2C presents the box plots of the AUC and ER values achieved by different models on all 12 ST samples. The ranks of all these models remain consistent across the two metrics, with LETSmix consistently holding the top position. Due to the greater sensitivity of the AUC metric compared to that of the ER metric, the evaluation results based on the AUC metric exhibit more pronounced differences in cell-type deconvolution performances across different models. Figure 2D shows the spatial distribution heatmaps of 3 layer-specific cell types estimated by each model trained exclusively on the “151673” ST sample. Compared to layer annotations, estimations of excitatory neurons from LETSmix were more accurate and coherent than those from other models. All excitatory neuron cell types estimated by LETSmix demonstrated reasonable regionally restricted patterns. For example, a distinct gap between layers 4 and 6 can be clearly observed in the LETSmix predictions for Ex_6_L4_6 cells. In contrast, the estimation results from other models were either excessively sparse or entirely fail to identify this cell type. Additionally, only LETSmix demonstrated the ability to correctly predict significantly more Ex_3_L4_5 cells within layers 4 and 5. Compared to CellDART, LETSmix exhibited a more continuous spatial distribution and fewer false positive results, crediting it to the utilization of information from spatial context. SpaDecon struggled to accurately predict the distribution of these cell types, mainly due to its neglect of the domain differences between the scRNA-seq and ST data during the modelling process. The same issue was also present in the CARD model. Although Cell2location also considered the domain shifts between data from the two sequencing technologies through traditional statistical probabilistic approaches, its performance fell short compared to that of the deep learning-based domain adaptation method employed in this study. Layer annotation information was also incorporated in POLARIS, but its estimation results evidently diffused into other non-target regions. Supplementary Figure S3 provides the results of all cell types estimated by LETSmix. Regarding other nonneuronal cells, LETSmix predicted that astrocytes are primarily distributed in layers 1 and 6, while oligodendrocytes were mainly located in the white matter region, which was consistent with findings from other biomedical studies (33), demonstrating the credibility and reliability of the predictions made by LETSmix.

Furthermore, LETSmix and other models were tested under three different conditions on the ST sample named “151673”. The AUC and ER values were calculated for each excitatory neuron type in 5 repeated experiments. As shown in Figure 2E, the scRNA-seq dataset was used with the original 28 cell types for model training under the “original” condition. These 28 cell types include several cell subtypes. In the “merge” condition, these subtypes were merged into a broader category before training the models. For example, Astros_1, Astros_2, and Astros_3 were merged into the Astros cell type. Only the 10 layer-specific excitatory neuronal cell subtypes used for metric calculations were not merged, resulting in 16 cell types in total. Under the “del Inhib” condition, all cells belonging to the inhibitory subtype, which accounts for approximately 20% of the entire dataset, were removed from the scRNA-seq data before model training. For both metrics and three conditions, LETSmix consistently achieved the highest scores, evidently outperforming other models, in accordance with the visual inspection in Figure 2D. To assess the robustness of different models under varying conditions, we visualized the predicted cell-type proportions of the ten neuronal cell types using UMAP representations (Figure 2F, Supplementary Figure S4A), and further quantified the centroid distances in UMAP space between estimation results across different condition pairs for each method (Figure 2G). Interestingly, the traditional machine learning-based methods (Cell2location, POLARIS, and CARD) demonstrated consistently higher stability when compared to deep learning-based approaches (LETSmix, CellDART, and SpaDecon). This enhanced stability may be attributed to the handcrafted features and fewer variable parameters in traditional machine learning models, which are less prone to overfitting under varying conditions. However, among the deep learning models, LETSmix exhibited smaller centroid distances overall, indicating better consistency across different conditions. Despite the superior stability of machine learning-based methods, they significantly lagged behind LETSmix in deconvolution accuracy. As a result, the apparent robustness of these traditional methods is of limited utility, as their lower prediction accuracy undermines their practical relevance in accurately resolving cell type compositions from spatial transcriptomics data.

In real-world applications, it is often challenging to obtain precisely annotated spatial regions from expert pathologists due to the scarcity or inaccessibility of such detailed annotations for tissue samples. However, with the rapid advancement of clustering methodologies applied to omics data, automated spatial region annotation has become increasingly feasible through the use of sophisticated computational tools (43-45). To explore this possibility, we evaluated the performance of LETSmix when employing clustering-based annotations, generated by GraphST (44), as a surrogate for expert-curated spatial regions in sample 151673 (Supplementary Figure S4B). The clustering-based annotations achieved an Adjusted Rand Index (ARI) score of 0.63 when compared to the ground truth annotations (Figure 4H), indicating a moderate level of agreement. To further assess the impact of this automated annotation approach on deconvolution performance, we compared results from LETSmix using the clustering-derived regions against the results obtained with expert annotations. Based on the AUC metric, no significant differences were observed between the two conditions (Figure 4I), supporting the feasibility of using clustering-derived annotations for deconvolution tasks. This finding underscores the potential of automated computational methods in addressing the limitations posed by the lack of expert annotations, especially in large-scale spatial transcriptomics studies. In addition to the clustering-based evaluation, we further tested performance of LETSmix when spatial contextual information and domain adaptation were systematically omitted. Both modifications led to a marked decline in performance, highlighting the critical importance of these components in maintaining the robustness and accuracy of the deconvolution process.

**Figure 3:**
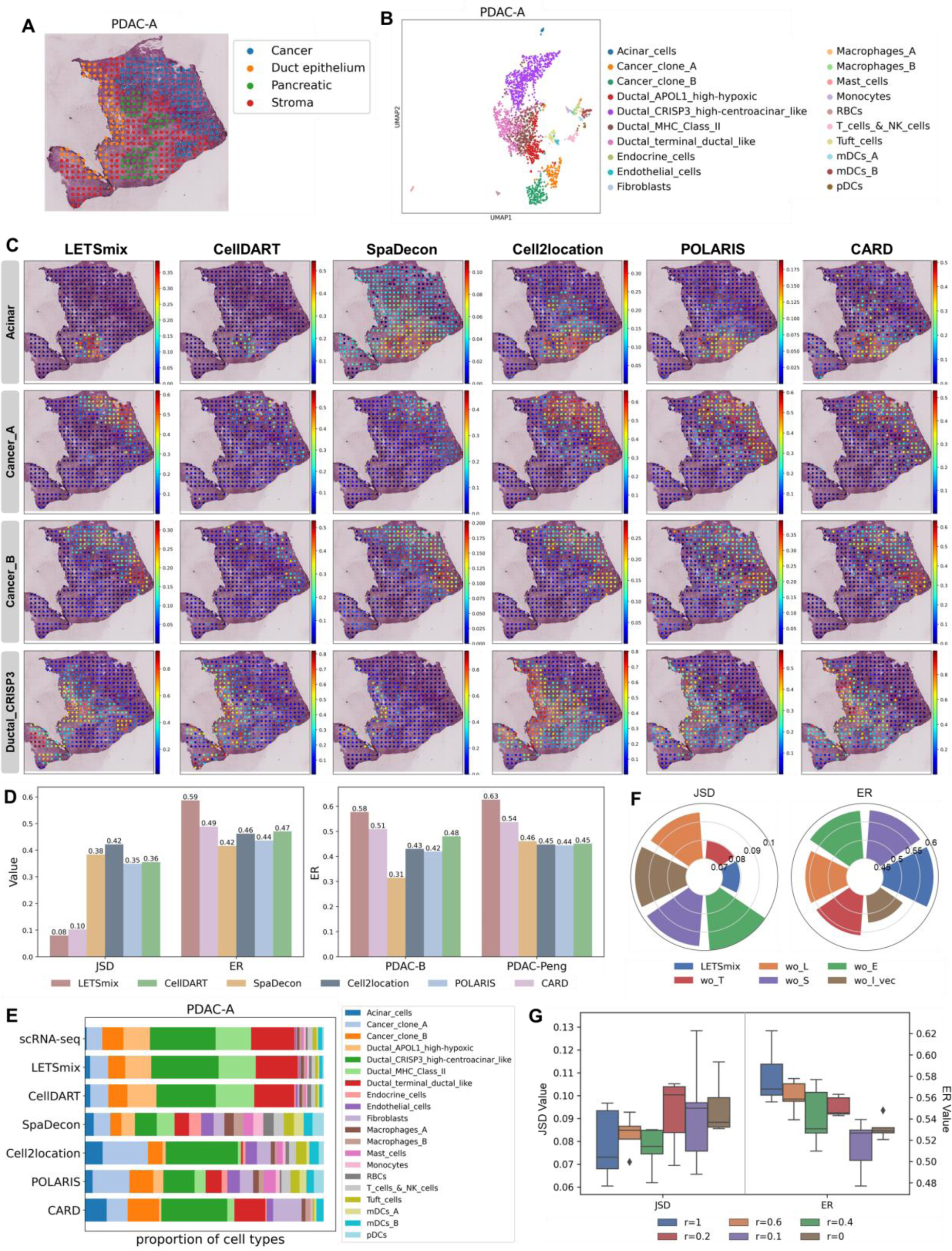
Application to the pancreatic ductal adenocarcinoma ST dataset. (**A**) Region annotations of the PDAC-A ST sample. (**B**) UMAP representation of the reference PDAC-A scRNA-seq dataset. (**C**) Estimated proportion heatmaps of 4 regionally restricted cell types by each model trained with matched PDAC-A ST and scRNA-seq data. (**D**) Left: model comparisons with matched ST and scRNA-seq data from PDAC-A. JSD and ER metrics were calculated using prior knowledge of cell type compositions and localizations, respectively, in PDAC-A tissue. Right: model comparisons through the ER metric evaluated in PDAC-A tissue, but models were trained with unmatched scRNA-seq data from PDAC-B and PDAC-Peng. (**E**) Stacked bar plots showing the overall cell type compositions in the PDAC-A ST sample estimated by each model using the paired PDAC-A scRNA-seq dataset. The ground truth was shown in the first row (denoted as “scRNA-seq”). The predicted proportion of each cell type is the average value of 5 repeated experiments. (**F**) Ablation study with the proposed LETS filter conducted on the matched PDAC-A dataset. “L”, “E”, “T”, “S” denote layer annotations, expression similarity, image texture features and spot coordinates, respectively. “l_vec” denotes the modified vectorized scaling factor *l*. (**G**) Performance of LETSmix using varying ratios of available ST spot data for the mixup-augmented domain adaptation training. The model was tested on the PDAC-A ST sample through ER and JSD metrics, and trained with matched scRNA-seq data. “r=0” denotes the situation without the mixup procedure. Each box plot ranges from the third and first quartiles with the median as the horizontal line, while whiskers represent 1.5 times the interquartile range from the lower and upper bounds of the box.

**Figure 4:**
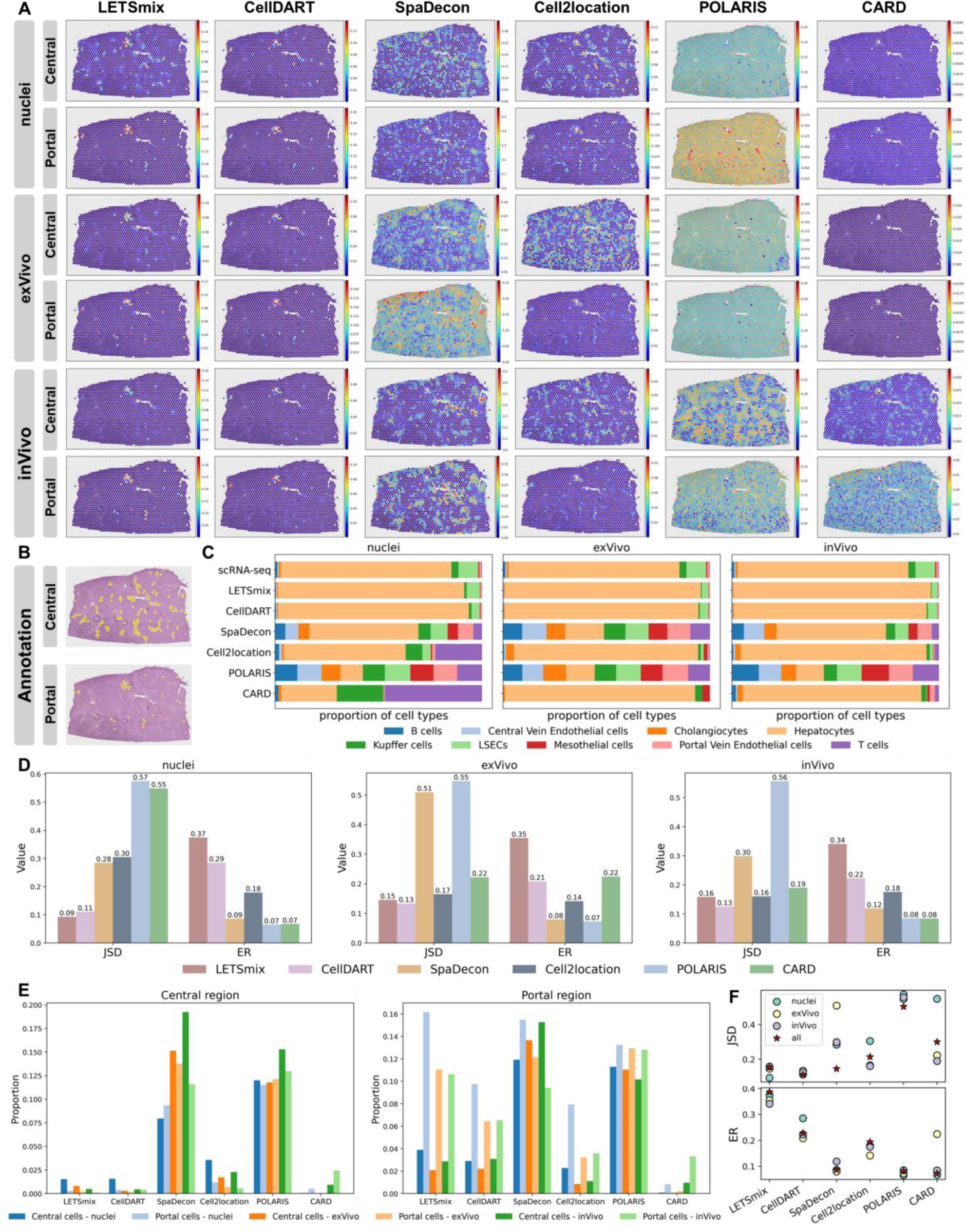
Application to the healthy mouse liver 10x Visium dataset. (A) Estimated proportion heatmaps of central vein and portal vein endothelial cells in the JBO001 Visium slice by each model trained with different reference scRNA-seq datasets. (B) Annotations of central and portal regions on the JBO001 ST sample. (C) Stacked bar plots showing the overall cell type compositions estimated by each model and the ground truth. The predicted proportion of each cell type is the average value in three Visium slices in 5 repeated experiments. (D) Model comparisons through JSD and ER metrics calculated using prior knowledge of cell type compositions and localizations, respectively, in mouse liver tissues. Each bar represents the average value of the involved cell types in three Visium slices and in 5 repeated experiments. (E) Comparisons of the average proportions of two cell types with regional distribution patterns in the target area estimated by each model. Each bar represents the average value of the involved cell type in three Visium slices and in 5 repeated experiments. (F) Scatter plots of metric values achieved by each deconvolution method under different experiment settings. “all” represents that the three scRNA-seq datasets are all used to train each model.

### LETSmix achieved superior and robust performance on PDAC data under matched and unmatched conditions

The second dataset used for evaluation originated from cancerous tissues from human PDAC patients. Here, ST and paired scRNA-seq data were collected following different protocols compared to those in the previously analyzed DLPFC dataset. We first applied LETSmix to an ST sample denoted as PDAC-A using paired scRNA-seq data for model training. The ST sample was delineated into four distinct regions by pathological experts (Figure 3A). Among the annotated cell types in scRNA-seq (Figure 3B), acinar, cancer, and ductal cells are expected to be located within specific regions (Supplementary Table S1). Fig 3C presents the spatial distribution pattern of cell types inferred by each model. Credited to the effective utilization of comprehensive spatial context information, acinar cells estimated by LETSmix were mainly distributed in the lower half of the pancreatic area, closely aligning with the manual annotation. In contrast, other models exhibited certain issues in inferring this cell type: some were excessively sparse (such as CellDART), while others generated numerous false-positive results, extending the predictions beyond the pancreatic area into stroma and cancer regions (as seen in SpaDecon and Cell2location). Cancer clone A and cancer clone B cell types are expected to be primarily distributed within the cancer region. The distinguishing marker genes for these two cell types are TM4SF1 and S100A4, respectively (Supplementary Figure S5A). TM4SF1 is significantly associated with tumor migration and invasion (46), indicating that areas with high TM4SF1 expression may represent late-stage cancer regions with metastatic potential. In contrast, S100A4 serves as an early prognostic marker for pancreatic cancer (47). Upon examination, the upper half of the cancer region in PDAC-A tissue exhibited increased TM4SF1 expression, while the lower half exhibited increased S100A4 expression (Supplementary Figure S5B). Thus, it was inferred that the upper region corresponds to late-stage cancer likely populated by cancer clone A cells, while the lower region represents early-stage cancer, possibly populated by cancer clone B cells. Among the results obtained from the five deconvolution models, LETSmix was able to most accurately identify such nuanced differences in the spatial distribution between the two cancer cell types. Although Cell2location generally delineated the cancer regions accurately, it failed to capture the relationship between these two cell types, incorrectly predicting a significant presence of Cancer clone A cells in the lower region. LETSmix also reasonably inferred the distributions of other cell types in PDAC-A tissue, consistent with the corresponding marker genes (Supplementary Figure S6). For example, spots enriched with TFF3, VIM, and CD74 marker genes were also estimated to have high proportions of ductal terminal, endothelial, and mDC cells, respectively.

Based on the test results under two quantification metrics in Figure 3D left, both the lowest JSD value and the highest ER value were achieved by LETSmix, showing that our proposed method was not only capable of mapping cells to their expected locations in ST, but also precisely estimating the proportions of various cell types. Similar to LETSmix, the CellDART model, which also employed domain adaptation techniques, outperformed other methods in the JSD metric evaluation, demonstrating the effectiveness of the applied adversarial training strategy in mitigating domain shifts between the two datasets. Although Cell2location performed well in predicting the enrichment of various cell types within specific regions, it ranked the poorest in the JSD metric, indicating its inability to accurately estimate the specific proportions of various cell types within each spot. This limitation was also reflected in the previous visualization of the prediction results for the two cancer subtypes. As visualized in Figure 3E, LETSmix and CellDART most accurately estimated the overall cell-type proportions in ST. Due to the complete neglection of domain differences, the input spot data exhibited significant distances from every clustering center in the feature space of SpaDecon trained on scRNA-seq data, leading to a similar estimated proportion for each cell type.

Next, the performance of different models was evaluated on scRNA-seq and ST data collected from another tissue region, denoted as PDAC-B (Supplementary Figure S7). Distributions of cancer, ductal centroacinar, and RBC cells predicted by each method are compared in Supplementary Figure S7C. Proportion heatmaps of the remaining cell types predicted by LETSmix are shown in Supplementary Figure S8, and Supplementary Figure S9 displays the distribution patterns of their corresponding marker genes for reference. Compared with the region annotations of the PDAC-B tissue shown in Supplementary Figure S7A, all models predicted the distribution of cancer clone A cells primarily within the cancer region. However, CARD incorrectly predicted a considerable number of cancer cells in the interstitium area, while SpaDecon estimated a very low percentage of cancer cells within the target region. Furthermore, the distribution of ductal centroacinar cells predicted by the SpaDecon model diffused from the ductal area to almost the entire tissue. As shown in Supplementary Figure S7D, similar to previous results on PDAC-A, although Cell2location and POLARIS performed well under the ER marker, their JSD values were significantly greater than those of the other models. In contrast, LETSmix achieved satisfactory performance in terms of both metrics. The presence of RBCs should be minimal or absent in pancreatic tissue, as PDAC tumors often compress and disrupt blood vessels, leading to reduced blood flow and impaired vascular function. This is also reflected in the stacked bar plot of the scRNA-seq data shown in Supplementary Figure S7E. However, except for LETSmix and CellDART, all other models estimated a substantial presence of RBCs in this tissue, which reaffirms the high accuracy of LETSmix in predicting the proportions of various cell types.

To further investigate the role of domain adaptation in the LETSmix model, we conducted additional tests in two scenarios of data mismatch. Specifically, when assessing cell-type deconvolution performance on the ST data from PDAC-A, models were trained using reference scRNA-seq data from PDAC-B (Supplementary Figure S10) and an external dataset denoted as PDAC-Peng (Supplementary Figure S11). The PDAC-Peng dataset was collected from 24 primary PDAC tumors and 11 normal pancreas tissues. Among the two ductal subtypes shown in Supplementary Figure S11A, ductal cell type 1 was identified as nonmalignant while the other was identified as malignant by previous studies (40), suggesting that ductal cell type 2 may infiltrate into the cancer region. According to the results obtained from the PDAC-B scRNA-seq data (Supplementary Figure S10C), predictions made by LETSmix were highly consistent with the previously obtained results using the matched PDAC-A scRNA-seq dataset. In contrast, other models exhibited significant changes in the predicted distributions compared to the previous outcomes. Specifically, acinar and ductal centroacinar cells estimated by other models exhibited severe diffusion into nontarget regions. Since scRNA-seq data from PDAC-B lack cells of the cancer clone B type, predictions for cancer clone A cells inferred by LETSmix appeared to be a fusion of both cancer cell types. Furthermore, except for LETSmix and Cell2location, other models mistakenly inferred a high presence of RBCs in this tissue. The visual inspection in Supplementary Figure S11C obtained by using data from PDAC-Peng further underscored that consistent prediction of acinar cells was made by LETSmix. Our proposed method also correctly estimated the distribution patterns of the two ductal subtypes, where non-malignant ductal cell type 1 was mainly enriched in the ductal region and malignant ductal cell type 2 was distributed across ductal and cancer regions. According to the ER metric shown in Figure 3D right, the superior performance of LETSmix over the other methods was even more pronounced than that achieved in Figure 3D left, which substantiated its stability in cell-type deconvolution when faced with considerable domain shifts between the scRNA-seq and ST data, demonstrating the ability to mitigate the need for alignment between these two data sources.

Building upon our preliminary investigation of the efficacy of the LETS filter in the DLPFC dataset (Figure 2I), we conducted a more comprehensive analysis by performing an ablation study to examine the contribution of each individual component of the LETS filter to the overall deconvolution performance (Figure 3F). Specifically, we systematically removed each element of the LETS filter—layer annotations, expression similarity, image texture features, and spot coordinates—and evaluated the subsequent effects on the deconvolution outcomes. The removal of image texture features resulted in the smallest decline in performance based on the JSD metric, yet it caused a significant reduction in the ER score, suggesting that while texture features may not drastically affect the global cell-type composition, they are essential for accurately identifying regionally enriched cell populations. Conversely, the removal of spot coordinates did not significantly affect the ER score, but it led to a pronounced increase in the JSD value, indicating that spatial information is crucial for maintaining overall consistency between the predicted and reference cell-type proportions. These findings highlight the complementary nature of the components within the LETS filter, as their combined use yields optimal deconvolution performance. Additionally, we observed that the modified vectorized scaling factor *l* also plays a critical role, as its removal led to a notable deterioration in both JSD and ER scores, underscoring its importance in balancing the integration of spatial and molecular features. Altogether, this ablation study demonstrates that individual components of the LETS filter contribute differently to the overall performance of LETSmix, and their synergistic integration is crucial for achieving the best deconvolution results.

Furthermore, given the limited number of available spots in the PDAC dataset, with only 428 spots in the PDAC-A sample and 224 spots in the PDAC-B sample, experiments were conducted to specifically assess the impact of the mixup data augmentation strategy integrated into the LETSmix model. We systematically evaluated the performance of LETSmix using different ratios of available spot data (Figure 3G). As expected, the deconvolution performance deteriorated progressively as the number of available spots decreased. Nevertheless, by applying the mixup augmentation, LETSmix was able to maintain performance even in the most extreme condition where only 10% of the spots were utilized, achieving results comparable to those obtained without mixup augmentation when the full dataset was used. This underscores the crucial role of mixup in maintaining model performance in scenarios with limited spatial transcriptomics data, such as those frequently encountered in clinical and experimental settings.

### LETSmix excelled in deconvolving complex spatial patterns in mouse liver using multiple scRNA-seq datasets

LETSmix was further applied to analyze three Visium slices of healthy mouse liver tissues (Supplementary Figure S12A). Three scRNA-seq datasets obtained with different experimental protocols, denoted nuclei, ex vivo, and in vivo, respectively, were used for joint analysis (Supplementary Figure S12B). Figure 4A illustrates the estimation results of two cell types with regional distribution patterns on the JBO001 Visium slice using three scRNA-seq datasets separately to train each model. Compared with ground truth region annotations shown in Figure 4B, it can be observed that LETSmix consistently provided the most accurate predictions among the tested methods for the spatial distribution patterns of these two cell types. However, it was also acknowledged that distributions of central vein ECs estimated by LETSmix slightly differed from that of the annotated central regions, and there was a tendency to misidentify substantial central vein ECs in the portal area. Yet, predictions for this cell type made by other methods also exhibited similar issues, potentially with more pronounced discrepancies. This difficulty may be attributed to the scarcity of central vein ECs within the applied scRNA-seq datasets (Figure 4C), limiting the ability of each model to sufficiently capture the characteristics of this cell type. Nevertheless, upon closer inspection of predicted central vein ECs by LETSmix (Supplementary Figure S12C), it can be observed that they still maintained a high similarity to the manually annotated central region, albeit with lower estimated proportions. Moreover, estimations for portal vein ECs by LETSmix were almost in perfect agreement with region annotations. LETSmix maintained high consistency in its estimation results when trained with three different scRNA-seq datasets. Cell2location also produced relatively decent estimation results for central vein ECs when trained with nuclei and in vivo scRNA-seq datasets, showing a certain correlation with the annotated central region. However, it predicted numerous false-positive results for both central vein ECs and portal vein ECs. Additionally, when ex vivo scRNA-seq was used as the reference dataset, Cell2location exhibited a significant decrease in deconvolution performance, indicating inferior stability compared to that of LETSmix. Although CellDART also produced stable prediction results with different scRNA-seq datasets, it estimated a relatively lower content for both types of ECs and did not accurately identify their locations compared to LETSmix. SpaDecon falsely predicted the occurrence of the two endothelial cell types across almost the entire tissue region, especially when trained with ex vivo scRNA-seq data. Similar issues were also observed in predictions made by POLARIS. In contrast, the CARD results hardly showed the presence of these two cell types. Although the estimation results for the two cell types generated by CARD trained on the in vivo dataset were also distributed throughout the entire tissue region, closer inspection revealed that the two predicted cell types accounted for only very small proportions, with the upper limit of the color bar much lower than 0.1. Furthermore, results given by SpaDecon and CARD exhibited significant differences when trained with different scRNA-seq datasets.

Quantitative evaluations further confirmed the exceptional performance of LETSmix compared to other deconvolution methods. Figure 4C visualizes the overall cell-type proportions across three ST Visium slices estimated by different models, and Figure 4D presents their JSD and ER scores. According to assessments based on the JSD metric, LETSmix and CellDART were the most accurate predictors for the general proportions of different cell types. Although CellDART achieved slightly lower JSD values using ex vivo and in vivo scRNA-seq data, LETSmix significantly outperformed CellDART in terms of the ER metric and maintained the highest accuracy in identifying the expected locations of each cell type across three scRNA-seq datasets. SpaDecon and POLARIS provided relatively uniform estimates for all cell types, which aligns with the experimental results observed in the PDAC dataset, with only hepatocytes being noticeably more abundant than other cell types in estimation results made by SpaDecon. On the other hand, when trained with the nuclei scRNA-seq dataset, Cell2location and CARD tended to predict an excessive number of T cells. Although this issue was alleviated in the ex vivo and in vivo results, where Cell2location and CARD achieved improved deconvolution performances, they were still significantly behind LETSmix. In fact, their performance on the PDAC dataset also surpassed that on the DLPFC dataset, with the former utilizing single-cell RNA-seq data and the latter utilizing single-nucleus RNA-seq data. This suggests a more pronounced domain shift between single-nucleus RNA-seq and ST data than between single-cell RNA-seq and ST data. Conversely, the performance of LETSmix on the nuclei scRNA-seq dataset was even slightly better than that on the ex vivo and in vivo datasets, as indicated by the lower JSD value and the higher ER value. A similar trend can be observed in the performance of CellDART, which also applies domain adaptation techniques. This implies that confounding information unrelated to the platform effect between ST and scRNA-seq data may be inadvertently introduced into features learned by the domain classifier when the degree of domain shift is inconspicuous. This, in turn, could impede the learning process of the source classifier.

Additionally, we investigated the differences between the estimated proportions of the two endothelial cell types within the central and portal regions, respectively, as shown in Figure 4E. In the central region, irrespective of the scRNA-seq dataset utilized, only LETSmix and Cell2location were able to accurately identify the quantitative relationship between the two cell types, with central vein ECs significantly outnumbering portal vein ECs. However, Cell2location also incorrectly estimated substantial portal vein ECs. CellDART achieved desirable results only when utilizing the nuclei scRNA-seq dataset. When trained with the other two scRNA-seq datasets, the proportions of both cell types predicted by CellDART were too low, and thus, their quantity differences were less distinct. SpaDecon, POLARIS and CARD exhibited suboptimal performance, producing unreasonable results where the number of portal vein ECs exceeded that of central vein ECs. Meanwhile, SpaDecon and POLARIS excessively estimated the proportions of the two cell types. In the portal region, LETSmix consistently predicted more portal vein ECs when trained with each scRNA-seq dataset. CellDART achieved similar results, but the predicted proportions of portal vein ECs were significantly lower than those of LETSmix, while the proportions of central vein ECs remained the same. Although SpaDecon and POLARIS predicted a large number of portal vein ECs in this region, they also inaccurately predicted a high proportion of central vein ECs.

Cell2location correctly identified a greater content of portal vein ECs in this area, but the disparity between portal and central vein ECs was less pronounced than that in LETSmix. CARD predicted excessively low content for both cell types when trained with the ex vivo scRNA-seq dataset, aligning with the observations in Figure 4A.

Finally, we investigated the performance differences when training the models using a combination of three scRNA-seq datasets compared to using each dataset individually (Figure 4F). While the use of multiple datasets can provide a more comprehensive representation of cellular heterogeneity and mitigate the risk of missing rare cell types due to insufficient data, it also introduces additional internal noise caused by batch effects. This added noise complicates the task of accurately learning cell-specific features, as the model must contend with variability between datasets. Our results show that only LETSmix and Cell2location demonstrated a slight improvement in ER values when trained on multiple datasets simultaneously. This improvement in Cell2location can be attributed to its explicit modelling of batch effects as a variable, which allows it to account for the discrepancies between datasets. LETSmix, on the other hand, was able to maintain performance by leveraging its spatial context integration and domain adaptation strategies, which help mitigate the impact of domain shifts across datasets. When assessing performance using the JSD metric, SpaDecon and POLARIS showed improved results when trained on all datasets concurrently. However, given the overall lower initial performance of these two models, the marginal improvements in JSD are of limited practical significance. Their initial poor performance suggests that despite the apparent gains in JSD, these models still struggle to provide accurate cell-type deconvolution, especially compared to LETSmix and Cell2location.

In summary, LETSmix predicts the spatial distribution of different cell types more accurately than CellDART, benefiting from the ability to utilize additional spatial context information in ST data. Although CARD also considers spatial correlations among spots in ST using their positional coordinates, the scattered distribution patterns of different regions within the Liver dataset make it challenging to accurately capture inherent correlations based solely on coordinate information. Similarly, POLARIS, which leverages region annotation information, struggles with this dataset due to the complex and irregular regional distribution, making it difficult to rely solely on such annotations for accurate deconvolution. In contrast, LETSmix overcomes these limitations and achieves superior performance by integrating multiple complementary sources of information. Credited to the use of domain adaptation techniques, only LETSmix and CellDART maintain high consistency in their estimation results when trained with the three different scRNA-seq datasets. This confirms that the proposed LETSmix model is more versatile and effectively alleviates the requirement for a high degree of matching between ST and scRNA-seq data.

### LETSmix demonstrated accurate cell-type deconvolution in single-cell resolution MOB data

With the continuous advancements in ST technologies, particularly in increasing spatial resolution, we sought to evaluate the performance of LETSmix on a MOB tissue ST dataset acquired by Stereo-seq, where spatial resolution reaches single-cell granularity. This dataset was divided into seven distinct anatomical layers, extending from the innermost to the outermost regions (Figure 5A). These regions were initially annotated on the DAPI-stained image, which, notably, lacked precise region labels for each individual spot. Based on prior analyses performed on the DLPFC dataset concerning the correspondence between ground truth region annotations and clustering results obtained from advanced computational methods (Figure 2I), the ConSpaS clustering model (45) was employed to infer the spatial region annotations for each spot in the MOB dataset (Figure 5B, Supplementary Figure S13A), providing a foundation for subsequent cell type deconvolution analysis. For the deconvolution task, we merged certain subtypes in the scRNA-seq dataset with similar UMAP distribution characteristics, reducing the original 38 cell types to a final set of 27 (Figure 5C, Supplementary Figure S13B). This refinement streamlined the analysis while maintaining sufficient granularity for distinguishing between biologically relevant cell types. Of particular interest in this study were cell types with potentially distinct spatial distribution patterns, as these could provide valuable insights into tissue organization and functional heterogeneity. To identify these cell types, we performed a correlation analysis between region-specific marker genes and the cell types in the scRNA-seq data (Figure 5D). Based on this analysis, we identified five cell types that displayed strong correlations with specific anatomical regions, showing strong potential for spatial enrichment (Supplementary Table S1), which were subsequently prioritized for focused analysis in the deconvolution task. Notably, unlike the previous datasets where H&E-stained images were utilized to construct the LETS filter for LETSmix, this MOB dataset provided single-channel DAPI-stained images, which primarily highlights the nuclei, providing a less comprehensive view of tissue morphology compared to H&E. Due to the smaller spot diameter in this ST dataset, the hyperparameter *k* in LETSmix was reduced from the default to 4, which controls the number of cells in each generated pseudo-spot.

**Figure 5:**
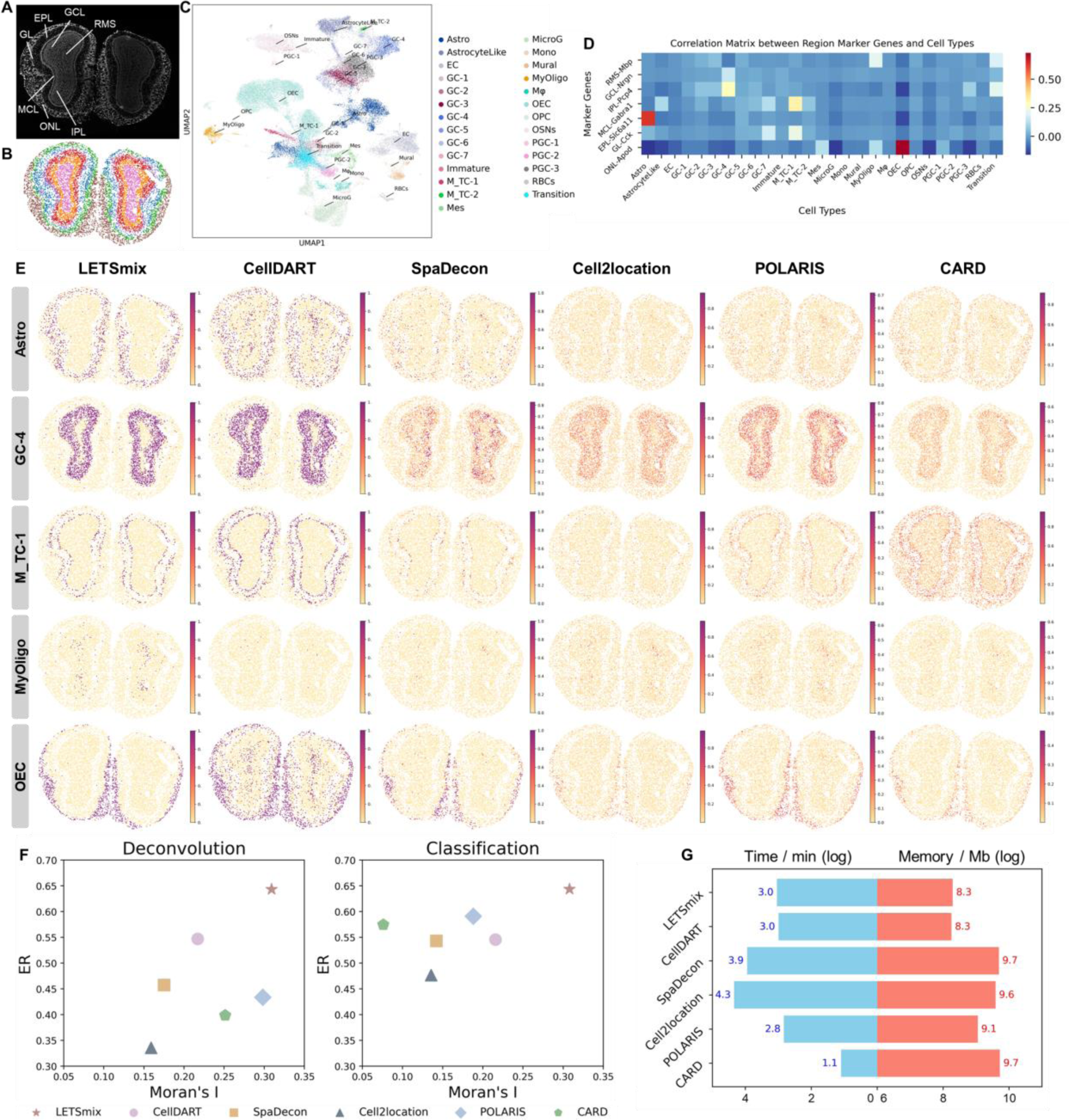
Application to the mouse olfactory bulb Stereo-seq dataset. (A) laminar structures of the MOB tissue annotated on the DAPI-stained image. (B) Spots clustering results generated by ConSpaS. (C) UMAP representation of the reference scRNA-seq dataset. (D) Correlation heatmap between region markers and cell types. (E) Proportion heatmaps of five cell types with potentially regional distribution patterns estimated by each model. (F) Scatter plots comparing the performance of different deconvolution models across two metrics. The left panel shows the results based on the predicted cell-type proportions for each spot. The right panel displays the performance after applying an argmax operation to assign each spot to the cell type with the highest predicted proportion. (G) Bar plots comparing the computational resource usage of each method based on their training time and peak memory usage.

The deconvolution performance of different models in predicting the spatial distribution of five key cell types with presumed spatial patterns is visualized in Figure 5E. LETSmix demonstrated superior accuracy in predicting the spatial distribution of these cell types, with results that were highly consistent with the distribution of known marker genes (Supplementary Figure S13C). Compared to other methods, LETSmix consistently showed higher sensitivity in capturing the distinct spatial regions associated with these cell types, as reflected by the more defined and concentrated patterns in the predicted distributions. This was particularly evident for GC-4 cells and OECs, where LETSmix predictions aligned with known anatomical knowledge, showing a clear enrichment in distinct regions of the MOB tissue. Astro cells predicted by LETSmix were predominantly located in the EPL region of the olfactory bulb, which was consistent with their known biological function in supporting synaptic transmission and maintaining the extracellular environment. In contrast, alternative methods such as SpaDecon, Cell2location, and CARD failed to clearly delineate the high prevalence of astro cells in this region, demonstrating a more diffused and less concentrated spatial distribution.

Considering that the ST data used in this study features single-cell resolution, it is reasonable to expect that each spot should predominantly represent a single cell type. This expectation was largely met by LETSmix and CellDART, where the predicted cell type distributions were characterized by deep, saturated colors, implying high confidence in the predicted cell types within each spot. The high confidence observed in their results also reflects the ability to accurately estimate the proportion of different cell types within each spot, which is consistent with their superior performance in previous experiments on the PDAC and Liver datasets, where both models achieved lower JSD values, further highlighting the benefits of incorporating domain adaptation. In contrast, other models, such as POLARIS, exhibited lower confidence in their deconvolution results. For instance, in the case of M_TC-1 cells, the upper limit of the color bar in predictions made by POLARIS reached only 0.6, indicating a lack of certainty in assigning this spot to specific cell types. This lower confidence may limit the ability to accurately capture the spatial distribution of cells, particularly in regions where sharp demarcations between cell types are expected.

To further explore the model predictions, we applied an argmax transformation to the deconvolution results, allowing us to identify the cell type with the highest predicted confidence for each spot. This transformation shifts the focus from cell-type proportion estimation (deconvolution) to spot classification. The classification results, visualized in Supplementary Figure S14, revealed notable changes in the predictions from SpaDecon, Cell2location, POLARIS, and CARD. In these models, the distribution of cells became more sparse following the argmax operation, likely reflecting low confidence in certain regions. Though CellDART and LETSmix both achieved robust performance, a more detailed comparison highlighted additional differences in their classification performance. From a biological standpoint, OECs are primarily involved in the support and regeneration of olfactory sensory neurons by ensheathing their axons. Due to their specialized role in olfactory neuron maintenance, they are typically confined to regions associated with olfactory nerve fibers and are not involved in the neuroblast migration that occurs in the RMS region. However, CellDART incorrectly predicted the presence of substantial OECs in the RMS, suggesting a misclassification in this area. Moreover, CellDART failed to identify the existence of MyOligo cells, further highlighting its limitations. By comparison, LETSmix not only avoided such errors but also demonstrated enhanced performance by incorporating spatial context into its predictions. This spatial awareness enables LETSmix to better capture the underlying biological processes and provide more accurate cell-type predictions across the tissue.

In addition to the visual inspection of the predicted cell-type distributions, we quantitatively evaluated the performance of different models using the ER and Moran’s I metrics under both cell-type deconvolution and spot classification tasks (Figure 5F). Consistent with previous observations, models other than LETSmix and CellDART exhibited significant differences in performance across these two tasks. While the transition from deconvolution to classification led to notable improvements in ER scores, a substantial decline in Moran’s I values was observed across these models, particularly for POLARIS and CARD. The increased ER values suggest that the argmax transformation effectively mitigates noise by reducing the presence of low-confidence predictions in non-target regions. Nonetheless, this transformation also led to a more scattered distribution of certain cell types, thereby decreasing the spatial coherence of the estimation results. In contrast, LETSmix consistently achieved significantly higher scores for both ER and Moran’s I, regardless of the task, demonstrating its robustness in preserving spatial coherence while achieving accurate cell-type predictions.

We also evaluated the computational resource consumption of each method on this dataset, focusing on both training time and memory usage (Figure 5G). LETSmix, despite incorporating multiple sources of information and leveraging sophisticated domain adaptation techniques, demonstrated moderate training time, placing it in the middle range among the methods tested. In terms of memory consumption, LETSmix exhibited a clear advantage due to its efficient code implementation and the LETS filter, which improves data quality and reduces the need for complex network architectures with numerous parameters. It is noteworthy that compared to CellDART, although LETSmix integrates additional spatial context information such as high-resolution histological images, as well as applying data augmentation strategies, the computational efficiency of the proposed method is nearly identical to that of CellDART. In summary, this balance between computational efficiency and predictive accuracy makes LETSmix a highly suitable choice for complex spatial transcriptomics applications.

## DISCUSSION

The identification of spatial distribution patterns for specific cell types plays a pivotal role in elucidating their positions, densities, and interactions within tissue structures, facilitating a comprehensive understanding of tissue complexity and pathological changes. Seq-based ST technologies measure average gene expression within cell mixtures. Through cell-type deconvolution, the positions and relative proportions of different cell types can be delineated on a spatial level, contributing to a more nuanced comprehension of tissue structure and cellular interactions. Additionally, spatial positions of spots and histological image information provide visual cues for tissue structure and cell distribution, enabling researchers to correlate ST data with specific cell types or structural features. This correlation aids in identifying differences and heterogeneity in cell types across different tissue regions, providing crucial insights for in-depth analysis. Furthermore, due to variations in data processing, detection sensitivity, technical specificity, cell handling, and sample preparation, certain domain shifts exist between ST and reference scRNA-seq data, which may impede the joint analysis of cell-type deconvolution. In this study, we introduce LETSmix, a deep learning-based method trained on pseudo-spots synthesized from reference scRNA-seq data, and real-spots augmented by mixing spots from ST data. LETSmix effectively utilizes spatial context information to construct a LEST filter, which enhances the continuity of the spatial distribution in deconvolution results and reduces noise in raw ST data. Moreover, LETSmix employs adversarial domain adaptation techniques to facilitate the seamless transition of robust deconvolution capabilities trained on simulated pseudo-ST to real ST data, enhancing the generalizability of LETSmix across different domains.

LETSmix excels in constructing precise spatial maps of cell type composition for ST samples. Evaluated across four datasets from distinct tissues, LETSmix notably outperforms other advanced deconvolution methods through both visual inspections and quantitative analyses leveraging prior knowledge of general cell-type locations and compositions. Beyond its effective deconvolution capability, LETSmix also boasts efficient computational resource consumption. Prior to model training, LETSmix conducts highly variable gene selection, significantly reducing the dimensionality of the input expression data. In the DLPFC dataset, a shared reference scRNA-seq dataset was used to deconvolve 12 ST samples. Marked domain differences exist between the scRNA-seq and ST data, and a high density of spots in ST indicates considerable structural features and noticeable stratification. These characteristics allowed the advantages of LETSmix to be fully manifested. In the visualization of layer-specific cell-type proportion heatmaps, predictions made by LETSmix closely aligned with layer annotations. Ablation and hyperparameter analyses conducted on this dataset strongly demonstrated the benefits of the employed domain adaptation technique and the utilized spatial context information. In the PDAC dataset, the performances of different models were evaluated using matched and unmatched scRNA-seq data as references to deconvolve two ST samples. Although fewer spots are available in this dataset and the domain shifts are less significant in the matched scenario, LETSmix still exhibits a nonnegligible performance advantage by mixing spots to effectively augment data samples in the real-ST domain. As the domain shift intensifies with unmatched reference scRNA-seq and ST data, LETSmix continued to reliably estimate cell-type distribution patterns, showing robustness. The performance of LETSmix was further assessed on mouse liver tissues using scRNA-seq data from different digestion protocols. Despite a weaker hierarchical structure among different functional regions, the incorporation of region annotation information enabled LETSmix to accurately capture inherent spatial correlations in ST data. Regardless of the scRNA-seq dataset used for model training, LETSmix consistently showed the ability to accurately determine the spatial distribution patterns of different cell types. In the final evaluation using the MOB dataset, which features spatial transcriptomics data at single-cell resolution and provides only single-channel DAPI-stained high-resolution images, LETSmix demonstrated its capability to accurately locate cell types with potentially regional distribution patterns by the use of clustering results generated from advanced computational tools. Predictions from LETSmix exhibited high confidence and robustness, highlighting the versatility of LETSmix in handling diverse ST datasets.

The LETS filter developed in this study serves as a versatile plug-in module designed to enhance the quality of ST data by capturing inherent spatial correlations. This filter can be seamlessly integrated into other deconvolution models, thereby expanding its utility beyond the LETSmix framework. By applying local smoothing to adjacent spots with similar morphological characteristics, the filter ensures that their corresponding expression profiles exhibit intended similarity, which in turn facilitates spatial continuity in deconvolution results. As clustering analysis in ST continues to advance, we demonstrated that state-of-the-art clustering models can be effectively utilized to guide the construction of the LETS filter, particularly in cases where manual region annotations are unavailable for the ST dataset. Notably, concerns may arise regarding the potential for the LETS filter to introduce “smoothing effects,” which could lead to the overshadowing of rare cell types by more abundant ones, thereby compromising the accuracy of rare cell type representation. However, our experimental results on the Liver dataset provide strong evidence to the contrary. Despite the high prevalence of hepatocytes, which dominate the tissue and reduce the relative abundance of other cell types, LETSmix was able to accurately capture the spatial distribution of rare cell types, such as central vein ECs, whose average proportion in spots was only approximately 1% (Figure 4C, E). These results demonstrate the robustness of LETSmix in detecting and preserving the characteristics of rare cell types.

To address the absence of actual cell-type proportion labels in real-ST datasets, LETSmix employs a pseudo-ST generation methodology similar to that of CellDART, where a fixed number of cells with random weights are selected from scRNA-seq data to synthesize the gene expression profile of each pseudo-spot. While this approach theoretically allows for the generation of a sufficient number of pseudo-spots to simulate various combinations of cell types, there remain opportunities for improvement in this synthesis process. Future research could investigate alternative strategies for refining the generation of pseudo-spots, particularly to tackle issues such as class-imbalanced sample sizes, in which the characteristics of rare cell types may be overshadowed by those of more dominant cell types. Furthermore, although the domain adaptation technique employed in LETSmix is enhanced by the mixup strategy, which helps mitigate the imbalance between source and target domains, it still relies on relatively conventional adversarial training approaches. These approaches may reveal limitations in certain application scenarios. For instance, when the degree of domain shift between ST and the reference scRNA-seq data is minimal, it becomes challenging for the domain discriminator in LETSmix to effectively distinguish between the source and target domains. In such cases, the learned confusing features may negatively influence the source classifier, resulting in performance that is potentially less optimal than when domain adaptation is entirely omitted. Additionally, the current domain adaptation strategy is primarily tailored for single-source, single-target domain applications. In scenarios where multiple scRNA-seq datasets are available as references for training the model, internal domain shifts between these multi-source domains may complicate the training process. Consequently, the use of multiple scRNA-seq datasets simultaneously may not necessarily lead to a significant improvement in deconvolution performance. We anticipate that future developments in domain adaptation algorithms will offer more suitable solutions for handling the complexities of ST data analysis.

In conclusion, LETSmix emerges as a valuable tool in the field of spatial transcriptomics, providing enhanced capabilities for cell-type deconvolution. Its incorporation of spatial context information and effective domain adaptation techniques contribute to its ability to accurately delineate spatial distribution patterns. Although LETSmix has already demonstrated superior performance compared to other state-of-the-art models across multiple datasets, there is still room for further improvements. We anticipate that the suggested method has broad applications in comprehensively mapping tissue architecture across diverse biological contexts, aiding biomedical researchers in understanding cellular interactions, developmental processes, and pathological mechanisms within complex biological systems.

## DATA AVAILABILITY

Four publicly available datasets were analyzed in this study. Raw data can be obtained through the following websites or GEO accession numbers: (1) Human DLPFC (ST: http://research.libd.org/spatialLIBD/; scRNA-seq: GSE144136); (2) Pancreatic Ductal Adenocarcinoma (ST: GSM3036911, GSM3405534; scRNA-seq: GSE111672, https://download.cncb.ac.cn/gsa/CRA001160/); (3) Mouse Liver (ST: GSM5764 414, GSM5764415, GSM5764416; scRNA-seq: GSE192740); and (4) Mouse olfactory bulb (ST: https://github.com/JinmiaoChenLab/SEDR_analyses; scRNA-seq: GSE121891. Additionally, we curated and made available all the aforementioned datasets, which can be downloaded directly from the following website: https://zenodo.org/records/11114959.

An open-source implementation of the LETSmix algorithm in Python is available at https://github.com/ZhanYangen/LETSmix/ or https://figshare.com/account/articles/27457419, including codes for data preprocessing, network construction, and model training.

## SUPPLEMENTARY DATA

Supplementary data are available at NAR online, including Supplementary Figures S1-14, Supplementary Tables S1-4, and Supplementary Algorithm S1.

## ACKNOWLEDGEMENTS

*Author contributions.* Yangen Zhan: Conceptualization, Investigation, Data curation, Methodology, Software, Formal Analysis, Visualization, Writing – original draft. Yongbing Zhang: Conceptualization, Funding acquisition, Project administration, Resources, Supervision, Writing – review & editing. Zheqi Hu: Data curation, Software, Validation. Yifeng Wang: Conceptualization, Writing – review & editing. Zirui Zhu: Data curation, Software, Validation. Sijing Du: Data curation, Software. Xiangming Yan: Software. Xiu Li: Funding acquisition, Project administration, Supervision, Writing – review & editing.

