## Supplementary Information for "LETSmix: a spatially informed and learning-based domain adaptation method for cell-type deconvolution in spatial transcriptomics"

### Supplementary Figures

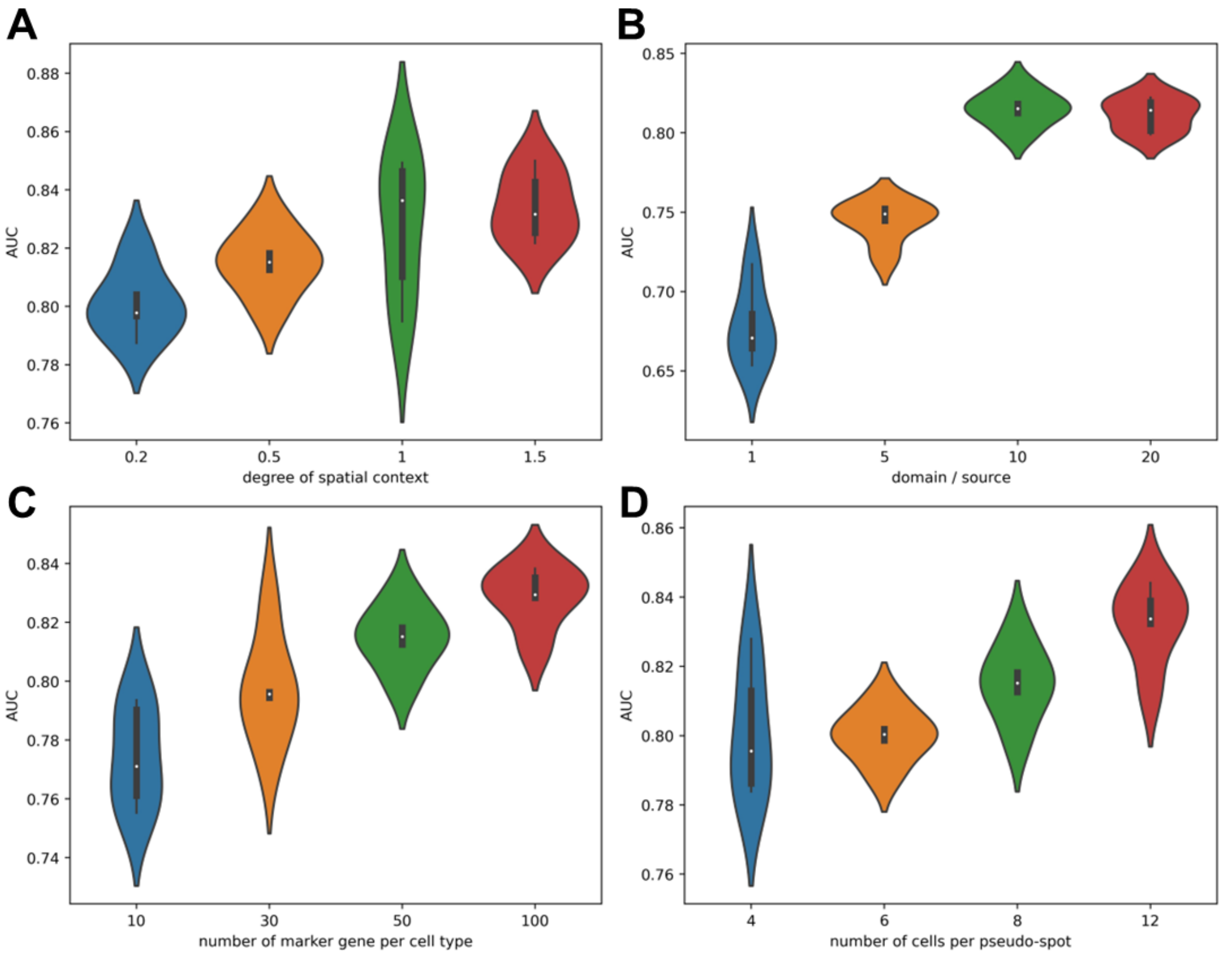


**Supplementary Figure S1: Experiments on different choices of 4 hyperparameters in LETSmix, evaluated on the DLPFC ST sample named "151673".** Each violin plot includes 5 datapoints, representing the mean AUC value for 10 layer-specific excitatory neuron cell types across five repeated experiments. **(A)** The hyperparameter "$\tilde{s}$" in the adjacent matrix construction process, representing the degree of spatial context information used. A higher value suggests a more extensive refinement of the ST dataset. (**B**) The hyperparameter "$d$" in the network adversarial training process, representing the number of iterations the domain classifier is trained after the source classifier in each cycle. **(C)** The hyperparameter "$m$" in the course of data preprocessing, representing the number of top marker genes selected for each cell type. **(D)** The hyperparameter "$k$" in the pseudo-spot generation process, representing the number of selected cells in each pseudo-spot.

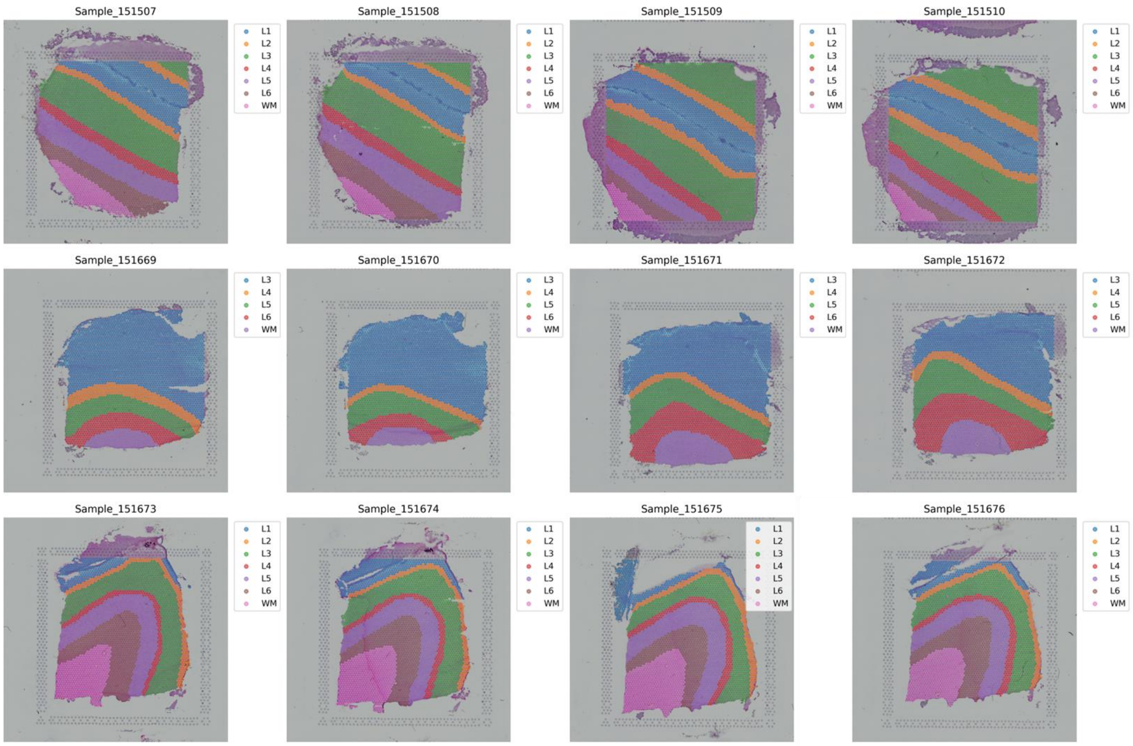


**Supplementary Figure S2: Layer annotations of all 12 DLPFC ST samples.**

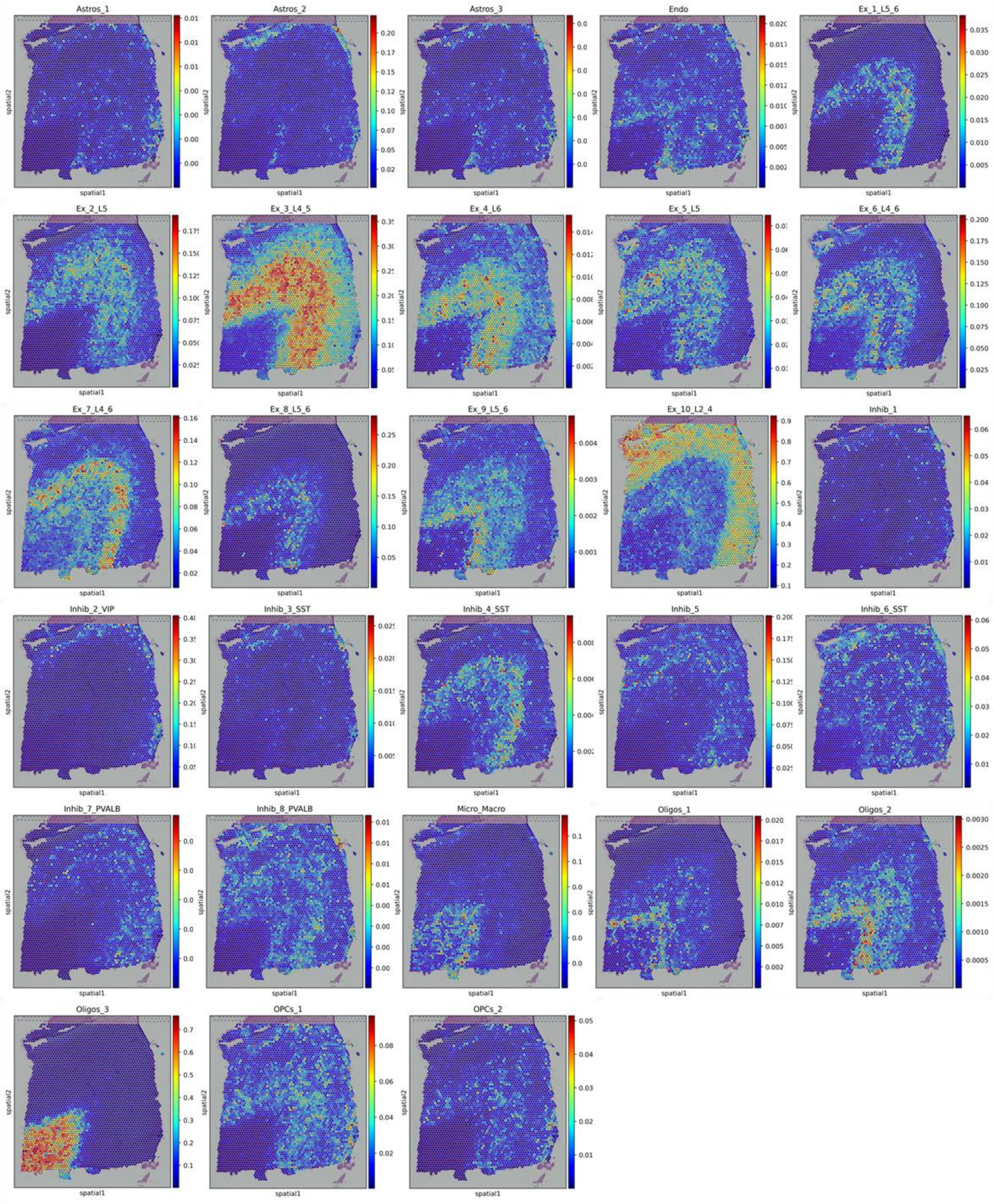


**Supplementary Figure S3: Proportion heatmaps of all 28 cell types in the 151673 ST sample estimated by LETSmix.**

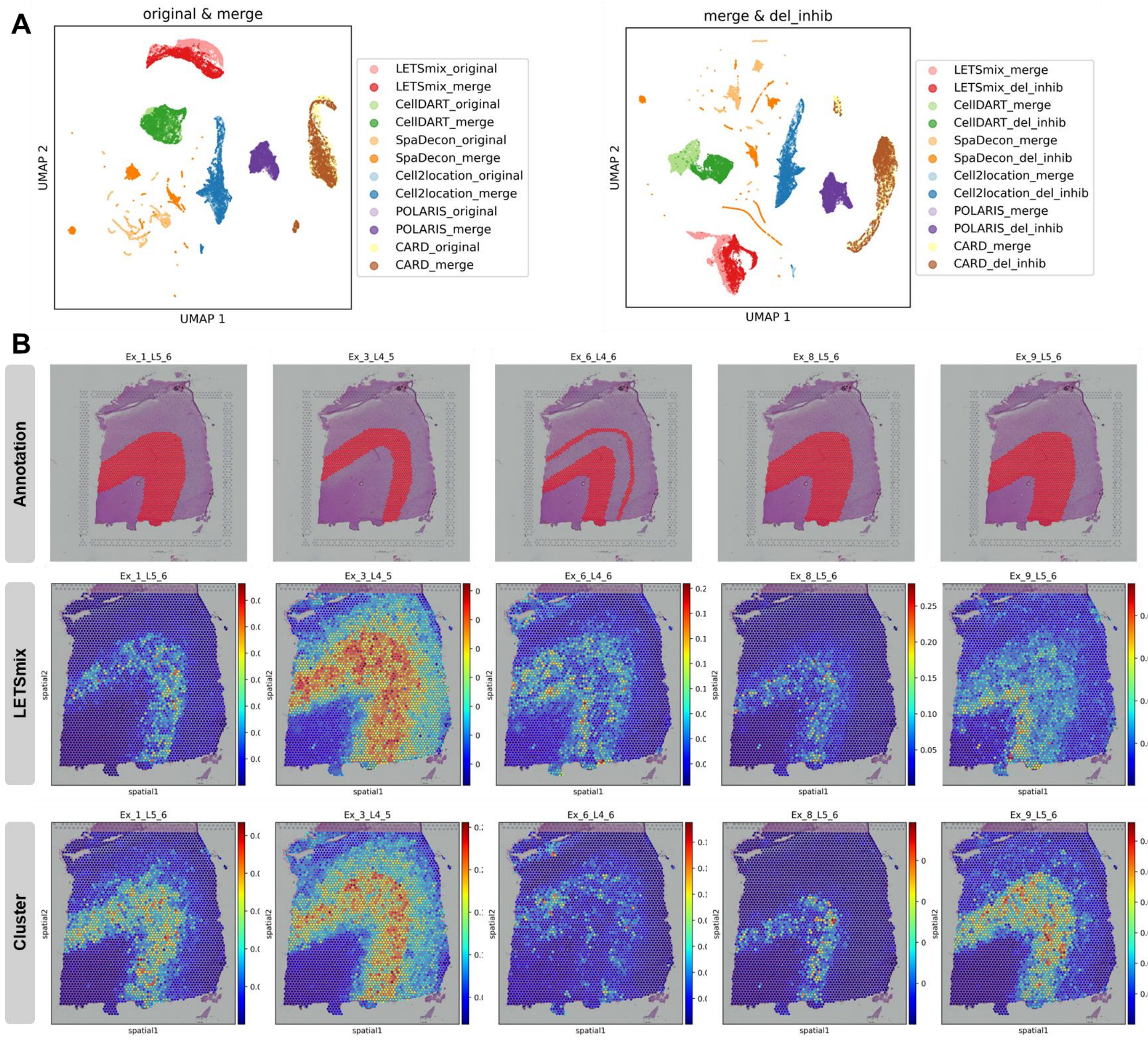


**Supplementary Figure S4: (A)** UMAP representation of deconvolution results from different methods under certain condition pairs. (**B)** Estimated proportion heatmaps of 5 layer-specific excitatory neurons by LETSmix. Results in the second row are obtained by using ground truth layer annotations, and results in the third row are obtained by using clustering results from GraphST.

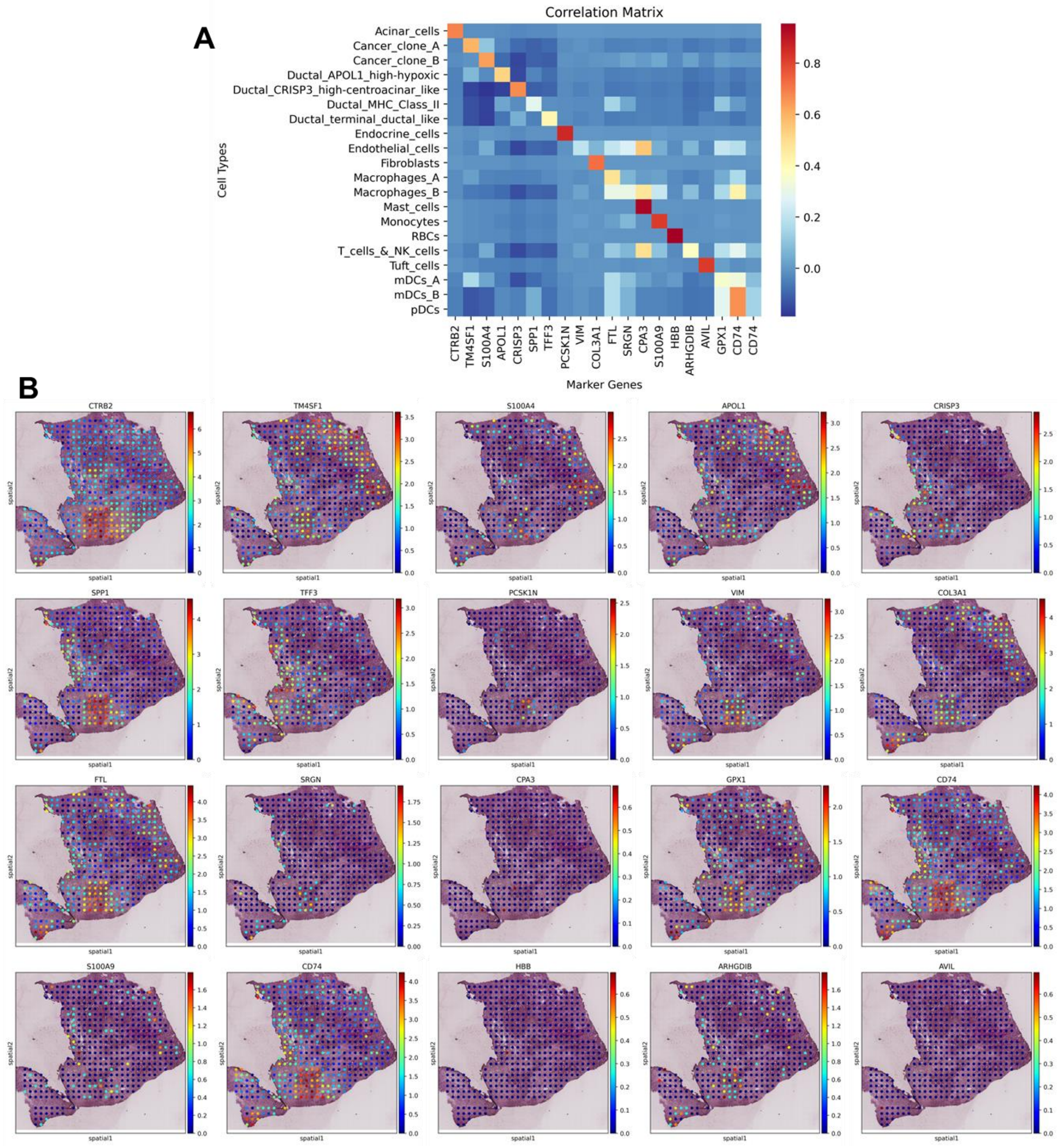


**Supplementary Figure S5: (A)** The correlation matrix for log-normalized gene expression in the top marker gene of each cell type in the PDAC-A scRNA-seq dataset. **(B)** Log-normalized gene expression heatmaps of these markers in the PDAC-A ST dataset.

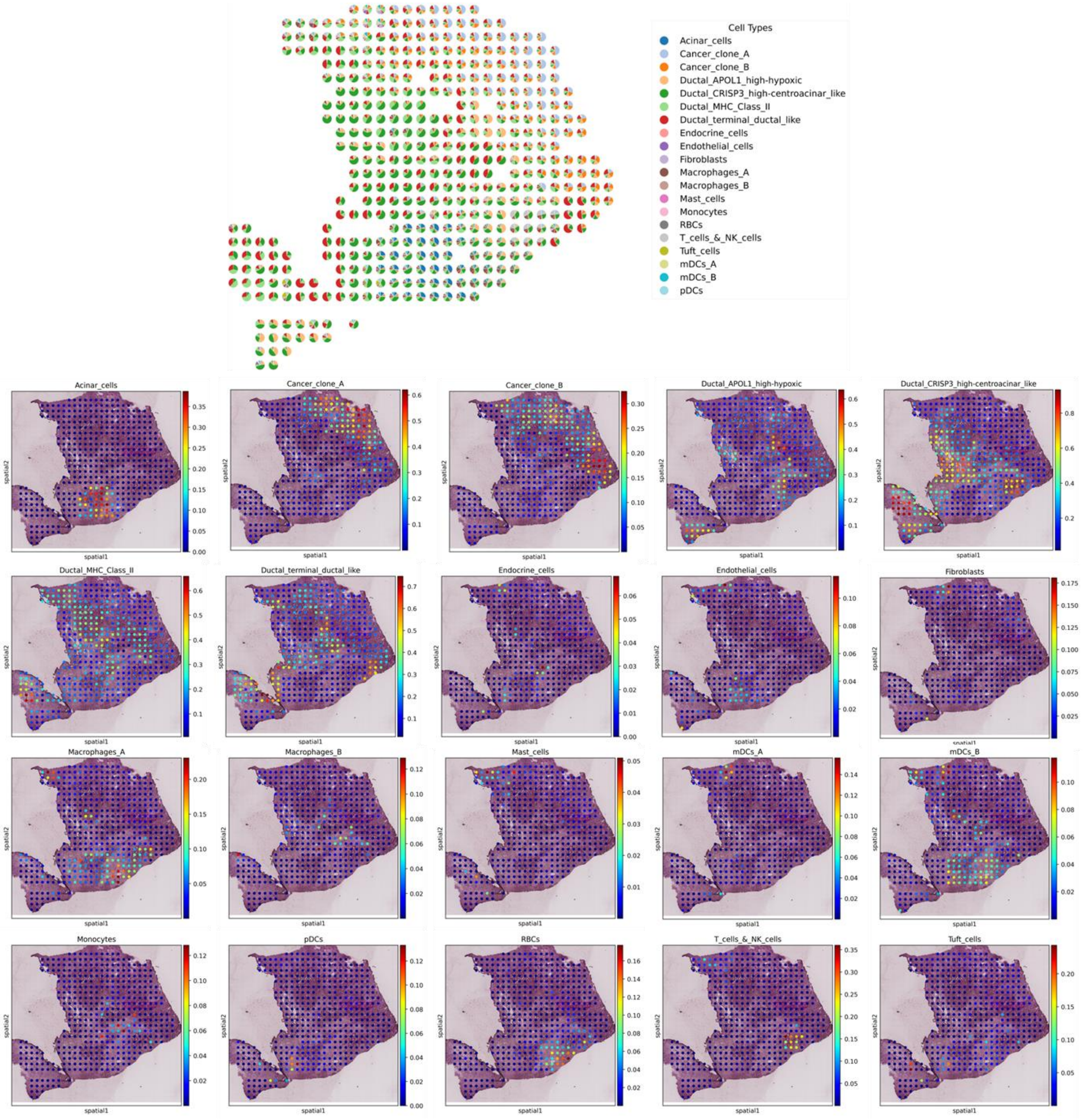


**Supplementary Figure S6: Estimated proportion heatmaps of all 20 cell types by LETSmix trained on matched PDAC-A scRNA-seq and ST datasets.**

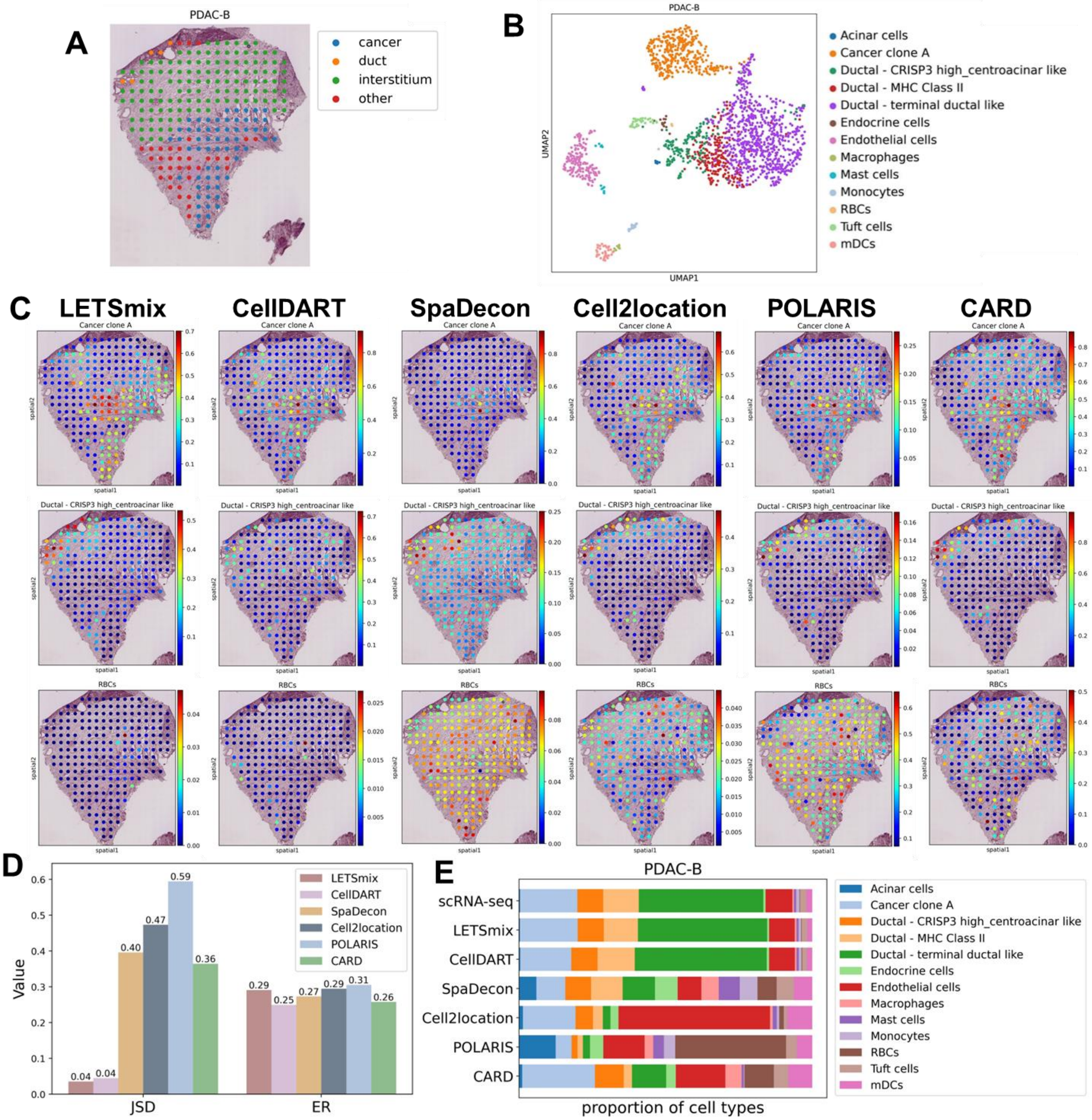


**Supplementary Figure S7: Application to pancreatic ductal adenocarcinoma ST dataset. Both scRNA-seq and ST data are from PDAC-B. (A)** Region annotations of the PDAC-B ST sample. **(B)** UMAP representation of the reference PDAC-B scRNA-seq dataset. **(C)** Estimated proportion heatmaps of 2 regionally restricted cell types and the RBCs that are expected to be rare in PDAC tissue. Ground truth region annotations are shown in the first column. **(D)** Model comparisons through JSD and ER metrics calculated using prior knowledge of cell-type compositions and localizations, respectively, in the PDAC-B tissue. Each bar represents the average value of the involved cell types in 5 repeated experiments. (**E**) Stacked bar plots showing the proportion of each cell type estimated by each model. The ground truth is shown in the first row (denoted as “scRNA-seq”). The predicted proportion of each cell type is the average value in 5 repeated experiments.

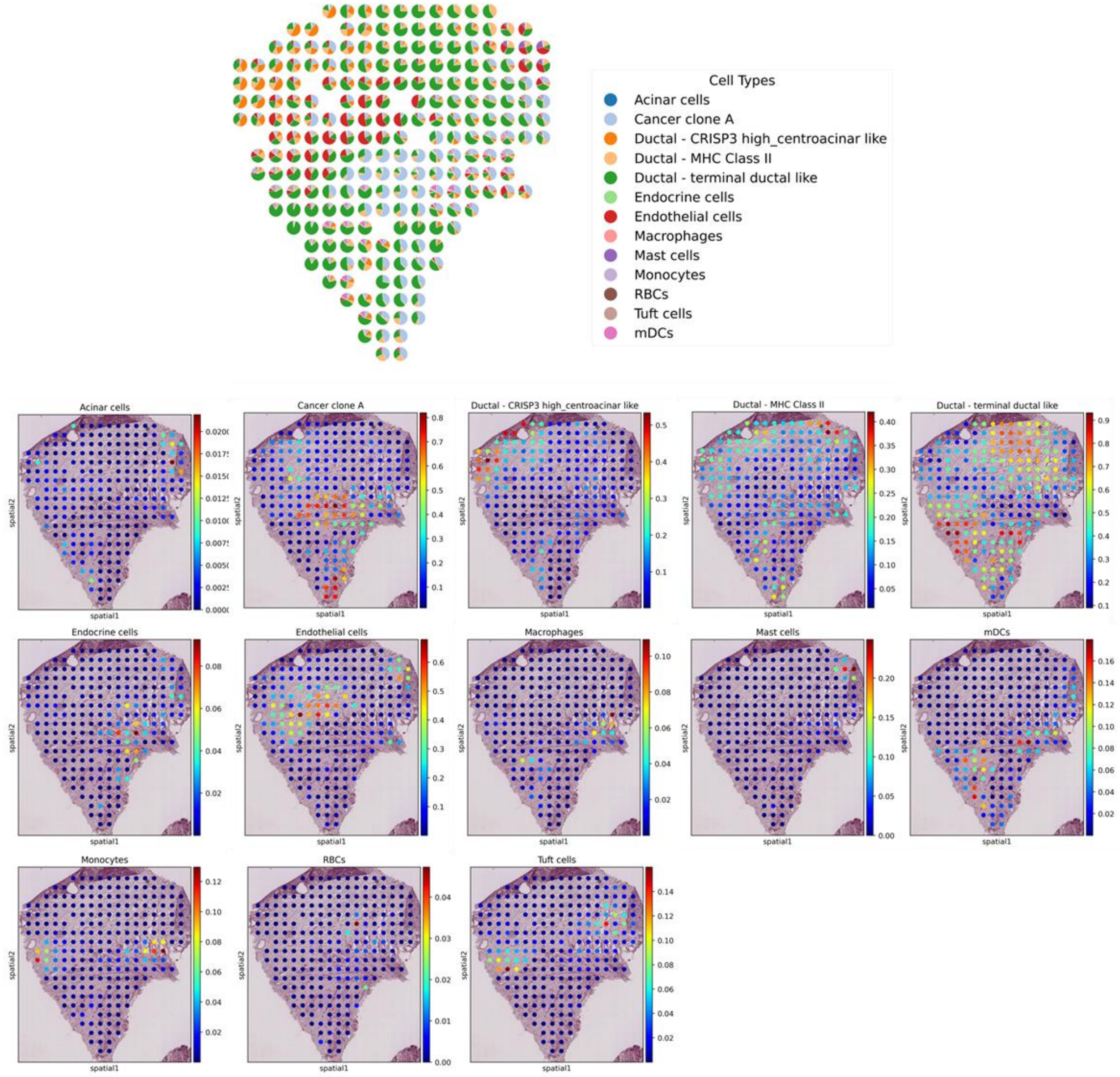


**Supplementary Figure S8: Estimated proportion heatmaps of all 13 cell types by LETSmix trained on matched PDAC-B scRNA-seq and ST datasets.**

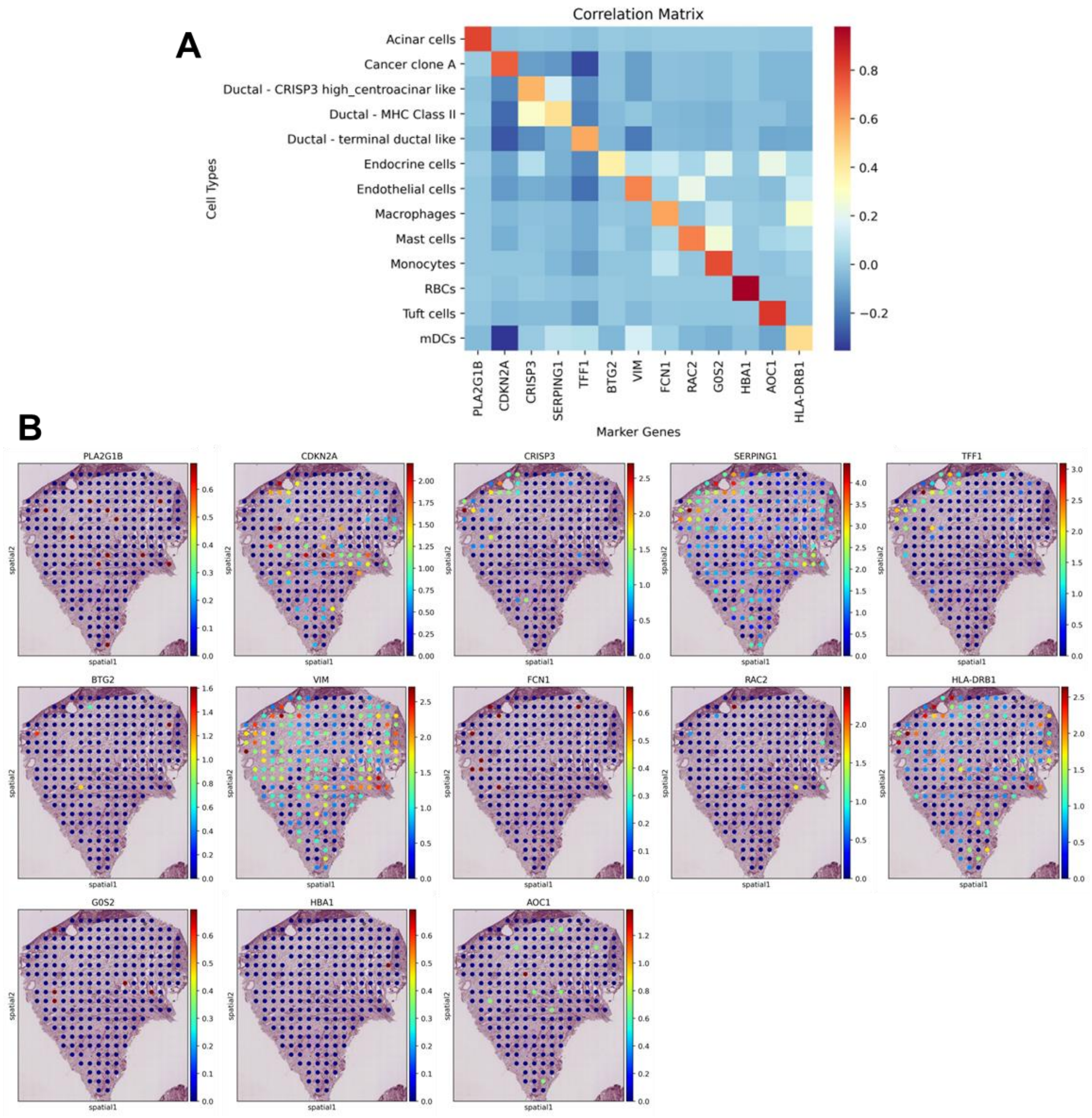


**Supplementary Figure S9: (A)** The correlation matrix for log-normalized gene expression in the top marker gene of each cell type in the PDAC-B scRNA-seq dataset. **(B)** Log-normalized gene expression heatmaps of these markers in the PDAC-B ST dataset.

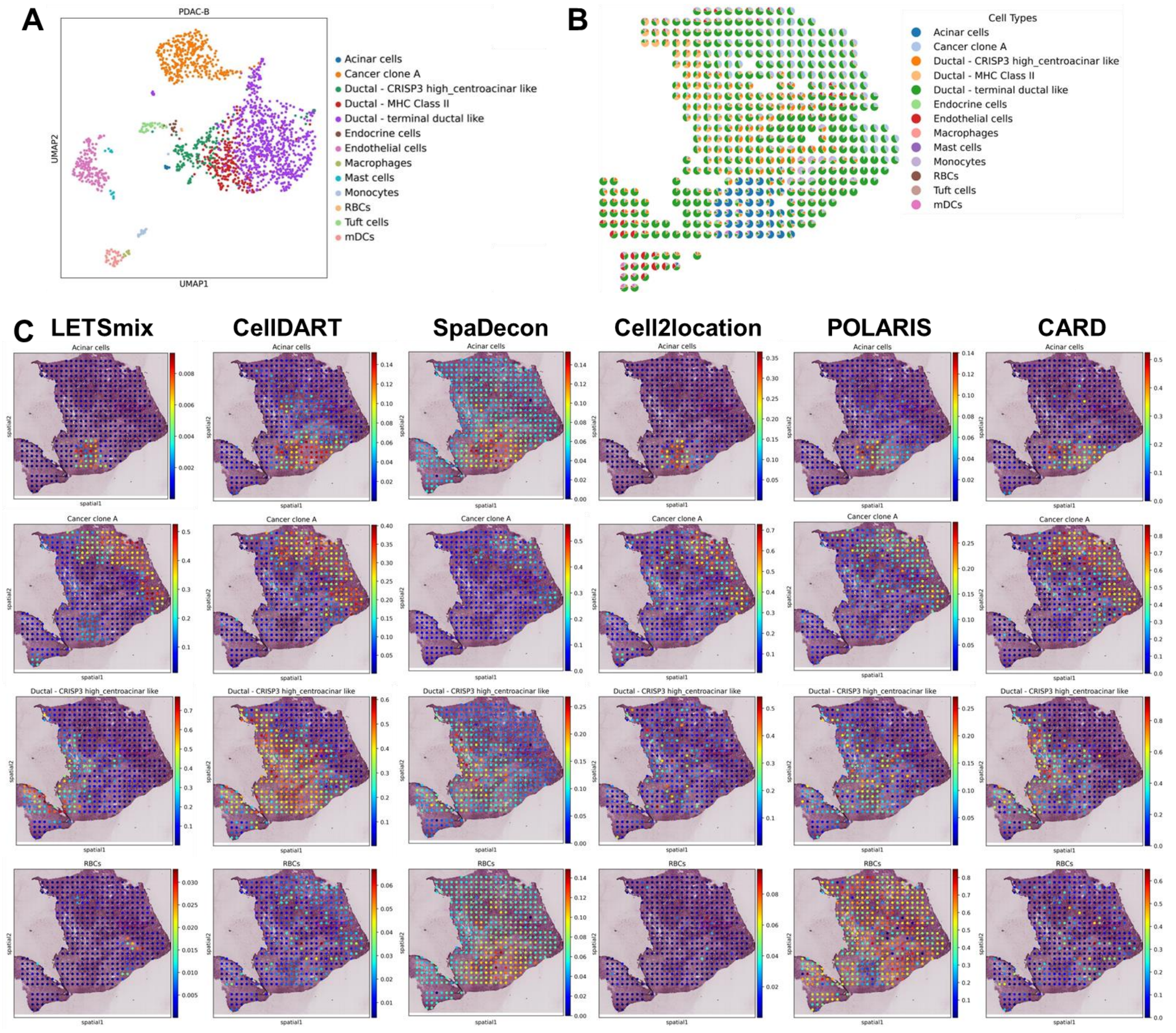


**Supplementary Figure S10: Application to pancreatic ductal adenocarcinoma ST dataset. scRNA-seq data are from PDAC-B while ST data are from PDAC-A. (A)** UMAP representation of the PDAC-B scRNA-seq dataset. **(B)** Pie plots display proportions and distribution patterns of all cell types estimated by LETSmix. **(C)** Estimated proportion heatmaps of 3 regionally restricted cell types and the RBCs that are expected to be rare in the PDAC tissue. Ground truth region annotations are shown in the first column.

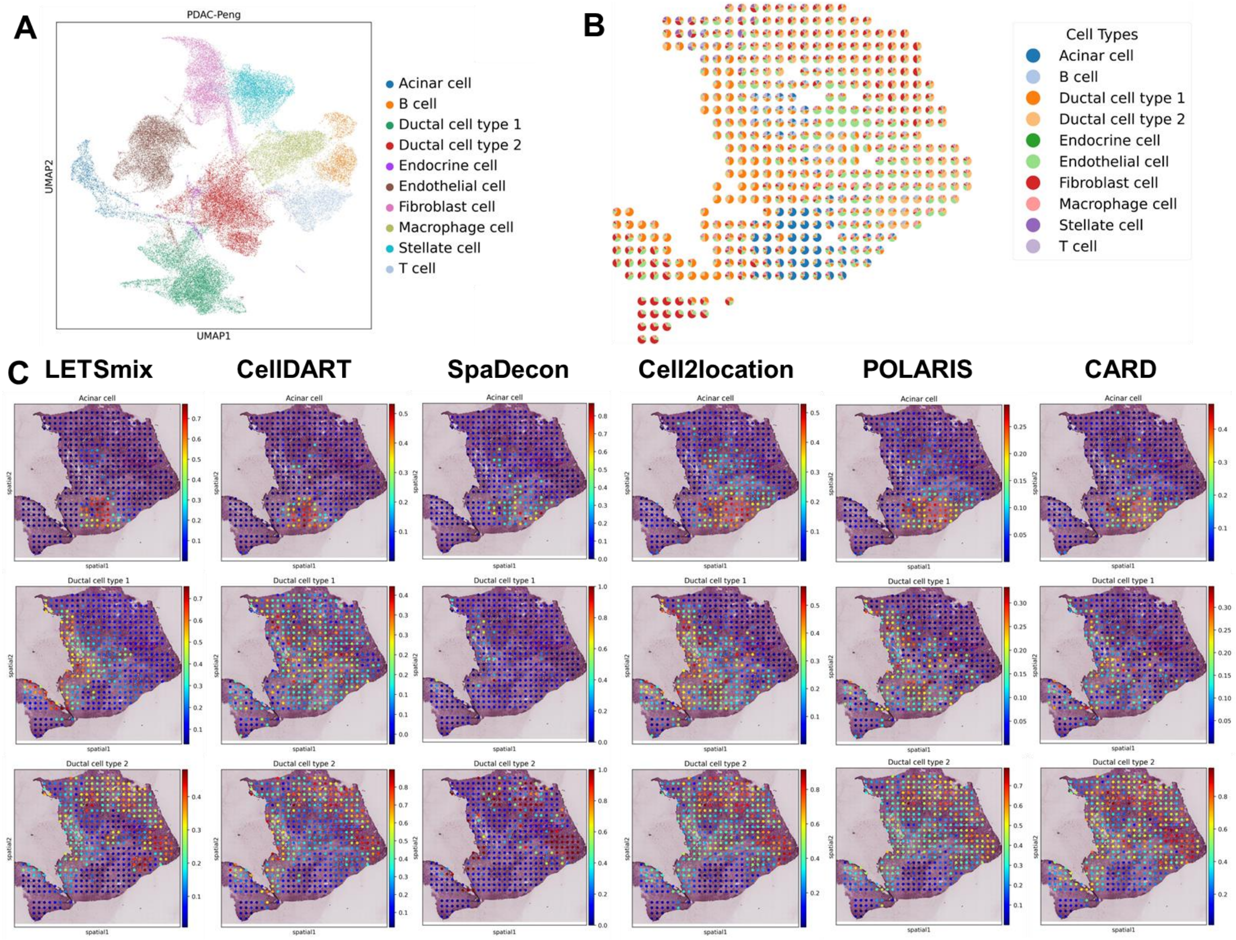


**Supplementary Figure S11: Application to pancreatic ductal adenocarcinoma ST dataset. scRNA-seq data are from PDAC-Peng while ST data are from PDAC-A. (A)** UMAP representation of the PDAC-Peng scRNA-seq dataset. **(B)** Pie plots display proportions and distribution patterns of all cell types estimated by LETSmix. **(C)** Estimated proportion heatmaps of 3 regionally restricted cell types. Ground truth region annotations are shown in the first column.

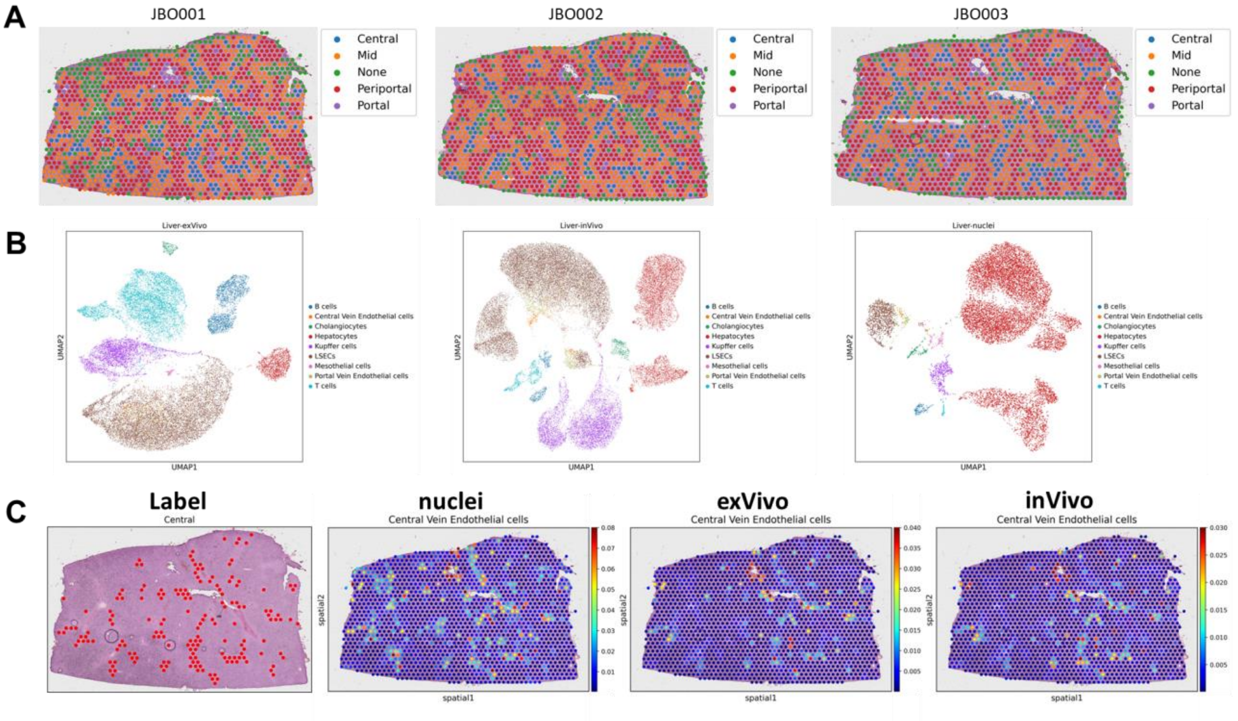


**Supplementary Figure S12: Visualization of the Liver dataset. (A)** Region annotations in three mouse liver sample slices. **(B)** UMAP representations of three scRNA-seq datasets with different experimental protocols. **(C)** Proportion heatmaps of central vein ECs estimated by LETSmix using three scRNA-seq datasets, respectively. Ground truth region annotations are shown in the first column. Compared to Figure 4A in the main text, the upper limit of the colorbar has been reduced to facilitate a clearer observation of the spatial distribution of this cell type.

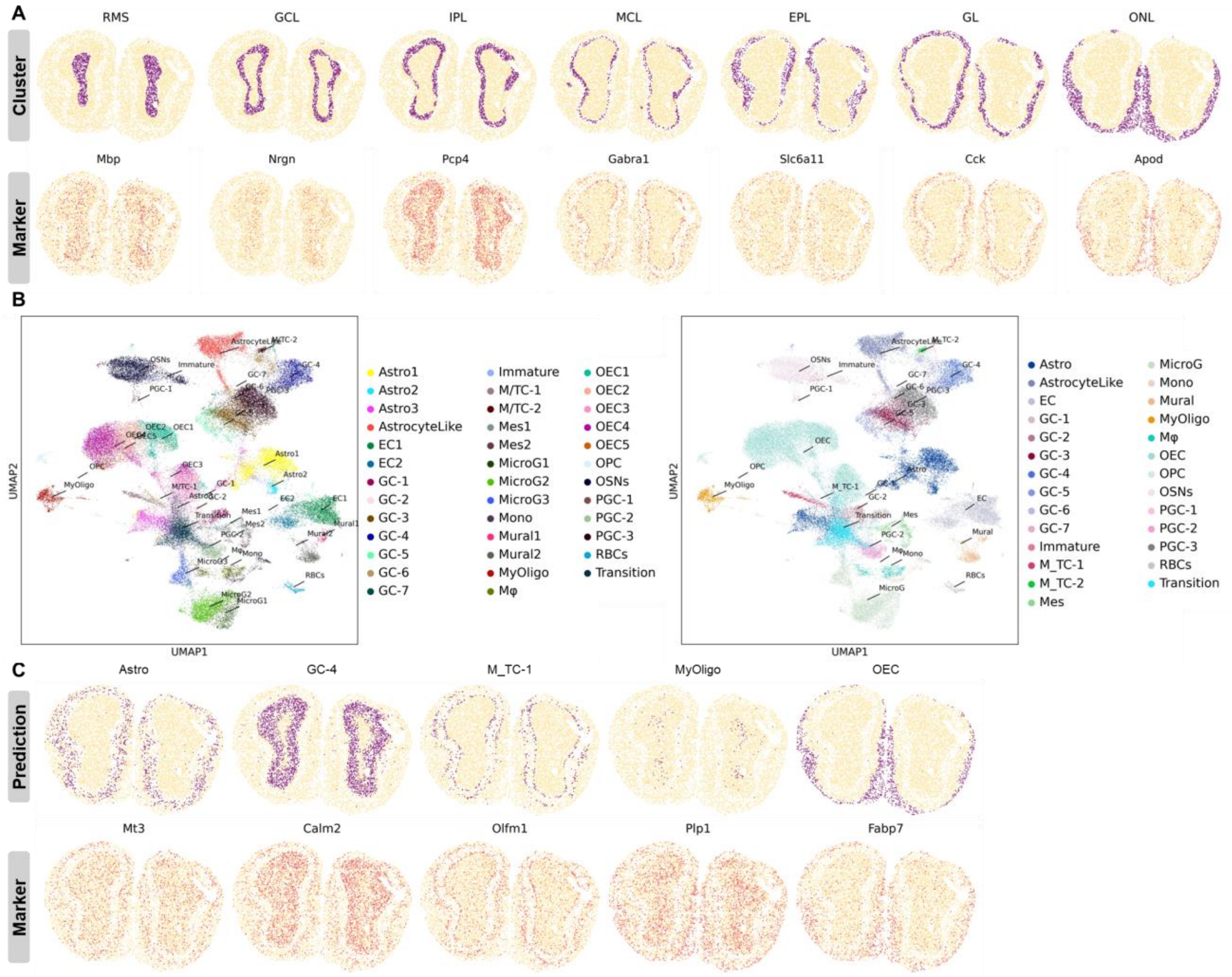


**Supplementary Figure S13: (A)** Clustering results generated by ConSpaS and the distribution of region maker genes. **(B)** UMAP representations of the original scRNA-seq dataset and the one after merging subtypes. **(C)** Cell type distributions estimated by LETSmix and distributions of the marker gene of each cell type.

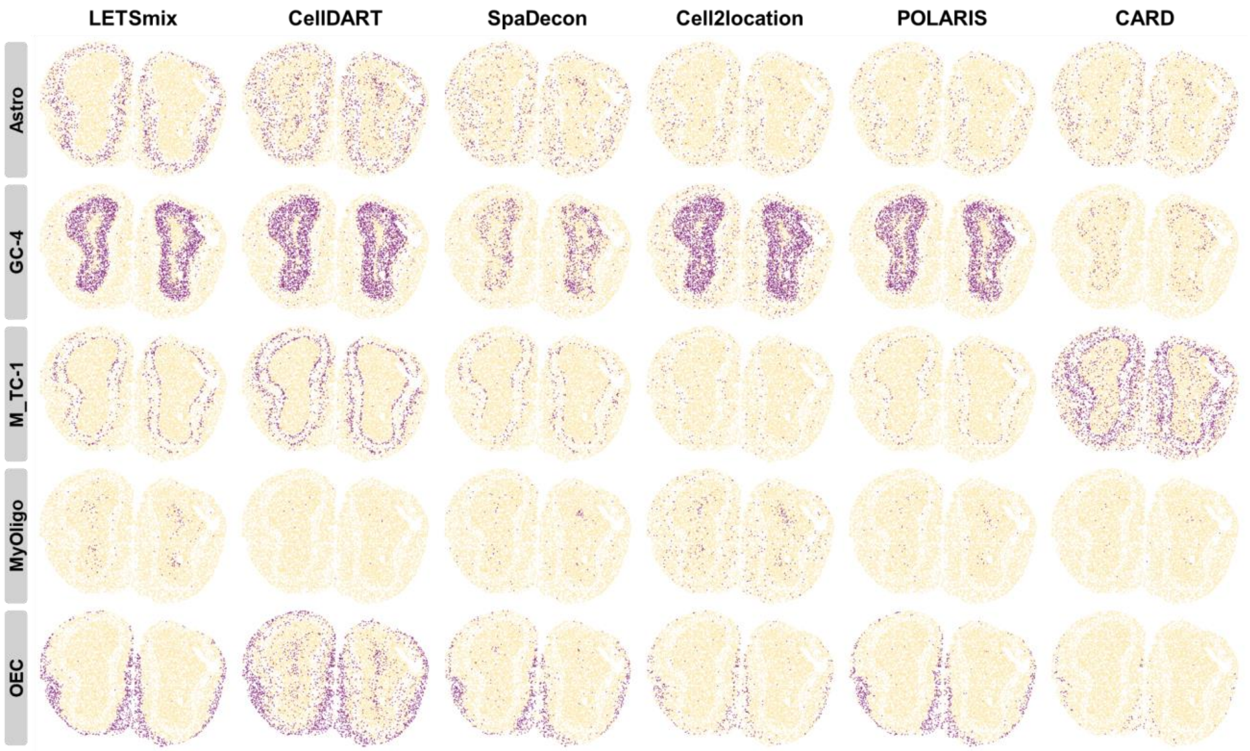


**Supplementary Figure S14: Distributions of cell types estimated by each method after argmax-transformed**.

### Supplementary Tables

**Supplementary Table S1: Layer-specific cell types in the scRNA-seq datasets used in this study and their expected enriched regions.**

| Datasets | Cell Types | Expected Regions | Datasets | Cell Types | Expected Regions |
|---|---|---|---|---|---|
| DLPFC | Ex_1_L5_6 | L5, L6 | PDAC-A | Cancer clone A | Cancer |
| | Ex_2_L5 | L5 | | Cancer clone B | Cancer |
| | Ex_3_L4_5 | L4, L5 | | Ductal - CRISP3 high centroacinar like | Duct epithelium |
| | Ex_4_L6 | L6 | | Ductal - MHC Class II | Duct epithelium |
| | Ex_5_L5 | L5 | | Ductal - terminal ductal like | Duct epithelium |
| | Ex_6_L4_6 | L4, L6 | | Acinar cell | Pancreatic |
| | Ex_7_L4_6 | L4, L6 | PDAC-B | Cancer clone A | Cancer |
| | Ex_8_L5_6 | L5, L6 | | Ductal - CRISP3 high centroacinar like | Duct epithelium |
| | Ex_9_L5_6 | L5, L6 | | Ductal - MHC Class II | Duct epithelium |
| | Ex_10_L2_4 | L2, L4 | | Ductal - terminal ductal like | Duct epithelium |
| PDAC -Peng | Acinar cell | Pancreatic | MOB | Astro | EPL |
| | Duct cell type 1 | Duct epithelium | | GC-4 | GCL, IPL |
| | Duct cell type 2 | Duct epithelium, Cancer | | M_TC-1 | MCL, GL |
| Liver | Central Vein Endothelial cell | Central | | MyOligo | RMS |
| | Portal Vein Endothelial cell | Portal | | OEC | ONL |

**Supplementary Table S2: Detailed network architecture in LETSmix**.

| Networks | Modules | dimensions |
|---|---|---|
| | Input expression data | n_gene |
| Feature extractor | Fully connected layer | 1024 |
| | Batch normalization | - |
| | ELU activation | - |
| | Fully connected layer | 64 |
| | Batch normalization | - |
| | ELU activation | - |
| Source classifier | Fully connected layer | n_cell_type |
| | Softmax activation | - |
| Domain classifier | Fully connected layer | 32 |
| | Batch normalization | - |
| | ELU activation | - |
| | Dropout (p=0.5) | - |
| | Fully connected layer | 2 |

**Supplementary Table S3: Datasets used in this study.**

| Datasets | Data Types | Technologies | Spots/Cells | Genes | Year |
|---|---|---|---|---|---|
| DLPFC | Spatial | 10x Visium | sample_151507: 4226<br>sample_151508: 4384<br>sample_151509: 4789<br>sample_151510: 4634<br>sample_151669: 3661<br>sample_151670: 3498<br>sample_151671: 4110<br>sample_151672: 4015<br>sample_151673: 3639<br>sample_151674: 3673<br>sample_151675: 3592<br>sample_151676: 3460 | 33538 | 2020 |
| PDAC-A | Spatial | Spatial Transcriptomics | 428 | 19738 | 2020 |
| PDAC-B | Spatial | Spatial Transcriptomics | 224 | 19738 | 2020 |
| Liver | Spatial | 10x Visium | sample_JBO001: 1646<br>sample_JBO002: 1651<br>sample_JBO003: 1604 | 31053 | 2022 |
| MOB | Spatial | Stereo-seq | 19109 | 27106 | 2022 |
| DLPFC | scRNA-seq | 10x Chromium | 56561 | 30062 | 2020 |
| PDAC-A | scRNA-seq | inDrop | 1926 | 19736 | 2020 |
| PDAC-B | scRNA-seq | inDrop | 1733 | 19736 | 2020 |
| PDAC-Peng | scRNA-seq | 10x Chromium | 57530 | 24005 | 2019 |
| Liver-nuclei | scRNA-seq | 10x Chromium | 12336 | 31053 | 2022 |
| Liver-InVivo | scRNA-seq | 10x Chromium | 34424 | 31053 | 2022 |
| Liver-ExVivo | scRNA-seq | 10x Chromium | 24512 | 31053 | 2022 |
| MOB | scRNA-seq | 10x Chromium | 51426 | 18560 | 2018 |

**Supplementary Table S4: Methods under comparison.**

| Algorithm | Paradigm | Spatial context | domain shifts | GPU | Languages | Strategies |
|---|---|---|---|---|---|---|
| LETSmix | Deep learning | √ | √ | √ | Python | Pseudo-spots with known cell type compositions are generated from reference scRNA-seq data, and are used to train the LETSmix model. Employing domain adaptation techniques ensures its deconvolution capability when applied to the real-ST dataset, which has been refined using spatial context information and augmented using mixup. |
| CellDART | Deep learning | - | √ | √ | Python | Similar to LETSmix, but without considering the potential spatial correlations among spots in the ST dataset. |
| SpaDecon | Deep learning | √ | - | - | Python | Similar to LETSmix, but without considering the domain shifts between scRNA-seq and ST data. It is trained in a supervised clustering manner with only the reference scRNA-seq data. |
| Cell2location | Statistical probabilistic | - | √ | √ | Python | Cell2location assumes that gene expression data in scRNA-seq and ST data follow a negative binomial distribution. Cell type signatures are inferred from the reference scRNA-seq dataset and used to estimate cell type proportions decomposed from ST data. The model incorporates parameters capturing various sources of data variability. |
| POLARIS | Statistical probabilistic | √ | - | √ | Python | Similar to Cell2location, it also adopts the negative binomial distribution assumption. While accounting for fewer sources of variability than Cell2location, it incorporates region annotations to allow for gene expression levels to vary across different tissue regions. |
| CARD | NMF regression | √ | - | - | R | The count matrics of scRNA-seq and ST data are decomposed into different parameter matrics. It infers cell type signatures from the reference scRNA-seq dataset and estimates the cell type composition matrix from ST data using NMF regression. The model introduces a conditional autoregressive assumption, leveraging position coordinates information to consider spatial correlations among spots. |

**Supplementary Algorithm S1**: Determine the value of the scaling factor $l = \{l_1, l_2, \cdots, l_N\}$ using an approximation method.

1) Set $l_i^0 = 1$ for $i \in \{1, 2, \cdots, N\}$, $\delta = 0.1$.
2) Calculate $s_i^\theta = \left(\sum_{j=1}^{N} a_{i,j}^\theta\right) - 1$,

   where $a_{i,j}^\theta = \exp\left(-\dfrac{d_{i,j}^2}{2l_i^\theta}\right) \times e_{i,j} \times m_{i,j}$, $\theta \in \{0, 1, 2, \cdots\}$.
3) $\begin{cases} l_i^{\theta+1} = l_i^\theta + \delta, \text{ if } s_i^\theta < \tilde{s} \\ l_i = l_i^\theta, \text{ otherwise} \end{cases}$
4) Repeat step (2) and step (3), until all values in $l$ have been determined.
